# Par3A and Par3B orchestrate podocyte architecture by regulating RhoA levels

**DOI:** 10.1101/2020.02.10.933671

**Authors:** Sybille Koehler, Johanna Odenthal, David Unnersjö Jess, Martin Höhne, Christian Jüngst, Ferdi Grawe, Martin Helmstädter, H. Henning Hagmann, Gerd Walz, Wilhelm Bloch, Carien Niessen, Bernhard Schermer, Andreas Wodarz, Barry Denholm, Thomas Benzing, Sandra Iden, Paul Thomas Brinkkoetter

**Author notes:** Corresponding author Paul Brinkkoetter Department II of Internal Medicine Center for Molecular Medicine Cologne (CMMC) University Hospital Cologne Kerpener Str. 62, 50937 Cologne, Germany.

## Abstract

Glomerular diseases are a major cause for chronic kidney disorders. In the majority of cases podocyte injury is causative for disease development. Cytoskeletal rearrangements and morphological changes are hallmark features of podocyte injury and result in dedifferentiation and subsequent loss of podocytes. Here, we establish a link between components of the Par3 polarity complex and actin regulators, which are necessary to establish and maintain the podocytes architecture utilizing both, mouse and *Drosophila* models. We demonstrate that the two mammalian Par3 proteins, Par3A and Par3B, share redundant functions despite differing in their ability to interact with other components of the Par complex. Only simultaneous inactivation of both Par3 proteins causes a severe disease phenotype in mouse podocytes by regulating Rho-GTP levels involving the actin regulators Synaptopodin and CD2AP in an aPKC independent manner.

## Introduction

The majority of kidney diseases originate from the glomeruli, small vascular units responsible for the filtration of plasma into a protein-free urine. A leaky filtration barrier and the presence of protein in the urine, i.e. proteinuria, are hallmarks of almost every glomerular disease and independently associated with an increased risk for end-stage renal disease and cardiovascular events ^1, 2^. The glomerular filtration barrier is composed of a fenestrated endothelium, the glomerular basement membrane (GBM) and podocytes. The latter are terminally differentiated, post-mitotic epithelial cells, which induce fenestrae formation within endothelial cells by secreting VEGF, maintain the GBM and act as outer cellular layer compressing the GBM and counteracting the filtration pressure to prevent the passage of macromolecules ^1, 3^. Podocytes exhibit primary and secondary foot processes and enwrap the glomerular capillaries completely. Foot processes from adjacent podocytes are joined by a specialized cell-cell contact known as the slit diaphragm (SD) ^1, 4^. During glomerular disease podocytes dedifferentiate into a columnar epithelial cell phenotype and plasma proteins leak into the urine. Ultimately, podocytes detach from the GBM and are lost into the urine leaving denuded capillaries resulting in glomerulosclerosis and progressive renal failure ^2^.

The importance of the podocytes’ architecture is emphasized by the fact that among the 50+ known gene defects leading to inherited forms of podocyte diseases are a large number of genes encoding for cytoskeleton-or SD-associated proteins ^5–7^. However, there is still limited knowledge about how podocytes are shaped. In single-layered, columnar epithelial cells, there are three major polarity complexes, i.e. the Par complex, the Scribble complex and the Crumbs complex. All three are essential to establish and maintain apico-basal polarity as well as for the formation of tight and adherens junctions^8^. Loss of Par3A or Scribble (SCRIB) did not result in glomerular phenotypes ^9, 10^. In contrast, loss of the atypical Protein Kinase C iota/lambda (aPKCiota), which is part of the Par complex, caused a severe glomerular disease phenotype ^11, 12^. Whilst these data rule out SCRIB as not being important for podocyte shape they are conflicting on the role of the Par complex.

Recently, we identified a second *Pard3* variant known as *Pard3B/Pard3L* ^13, 14^ to be expressed in podocytes. Par3B protein function remains elusive, in particular, whether the two Par3 proteins might have compensatory functions. Par3B shows high sequence similarities to Par3A with respect to its three PDZ domains and localizes to podocyte foot processes similar to Par3A ^9^. In contrast to Par3A, Par3B does not contain a classical aPKC binding domain and, as a consequence, does not directly interact with aPKC ^13, 14^. Here, we utilized the *Drosophila* nephrocyte model, as they serve as homolog cells to mammalian podocytes ^15^. Nephrocytes share evolutionary conserved functional similarities with mammalian podocytes as they filter fly hemolymph and also present with a highly conserved morphology including foot processes connected by the nephrocyte diaphragm ^15, 16^. This highly specialized cell contact is build out of homolog proteins to the mammalian slit diaphragm proteins Nephrin (*Drosophila Sticks-and-stones Sns*) and Neph1 (*Drosophila Dumbfounded Duf*) ^15, 17^. We show a critical role for Par3 proteins in controlling podocyte architecture. Par3A and B have redundant functions, as only loss of both orthologues resulted in a severe glomerular disease phenotype. We demonstrate, that the role of the Par complex is highly conserved, as loss of Baz, the only Par homolog in *Drosophila melanogaster*, results in loss of shape and function in the nephrocyte as well. Rho-GTP serves as downstream effector of the Par complex involving the actin-associated proteins CD2AP and Synaptopodin. Taken together, we identified how Par3A and Par3B link elements of polarity signaling and actin regulators to maintain podocytes’ architecture.

## Results

### Simultaneous loss of Par3A and Par3B causes a severe glomerular disease phenotype

In contrast to *Pard3a*, which is expressed ubiquitously throughout the whole glomerulus, *Pard3b* is expressed specifically in podocytes, where the protein localizes to foot processes (Supp. Figure 1) ^9^. This suggests a putative role for Par3B in the podocyte. Further, co-expression of Par3A and Par3B in podocytes raise the possibility of functional redundancy, which may explain the lack of phenotype observed for mutations in *Pard3a* alone. To investigate Par3A/B function further, we employed conditional *in vivo* targeting strategies specifically in podocytes to delete *Pard3b* alone or both *Pard3a* and *Pard3b* together. Podocyte specific gene deletion of *Pard3b* did not result in an overt glomerular disease phenotype, either in mice held on a pure C57/Bl6 or in a more susceptible CD-1 background (Figure 1, Supp. Figure 2+3). This is in line with the lack of phenotype in podocyte specific deletion of *Pard3a* ^9^. Although Par3A and Par3B single knockout mice did not present with proteinuria or severe foot process effacement, high-resolution microscopy revealed mildly effaced foot processes compared to control mice. This difference was significant as quantified by measuring the slit diaphragm length per capillary area. (Supp. Figure 4D). To study a functional redundancy between the two Par3 proteins, we generated podocyte-specific *Pard3a/b* double knockout mice (pure CD-1 background). Loss of both *Pard3* genes resulted in a severe glomerular disease phenotype with significantly increased urinary albumin levels already evident after 6 weeks (Figure 1A). Histological analyses revealed loss of the podocytes’ ultrastructure including foot process effacement (Figure 1B,E, Supp. Figure 4C,D), glomerulosclerosis and protein casts in the tubular apparatus (Figure 1C,D). These data show Par3 activity is required in podocytes to maintain their morphology and function. They further reveal that Par3A and B have redundant functions in podocytes. It is intriguing that Par3B is able to compensate for the Par3A loss despite Par3B inability to interact with aPKC. This suggests that Par3 function in the podocyte is independent of aPKC.

**Figure 1:**
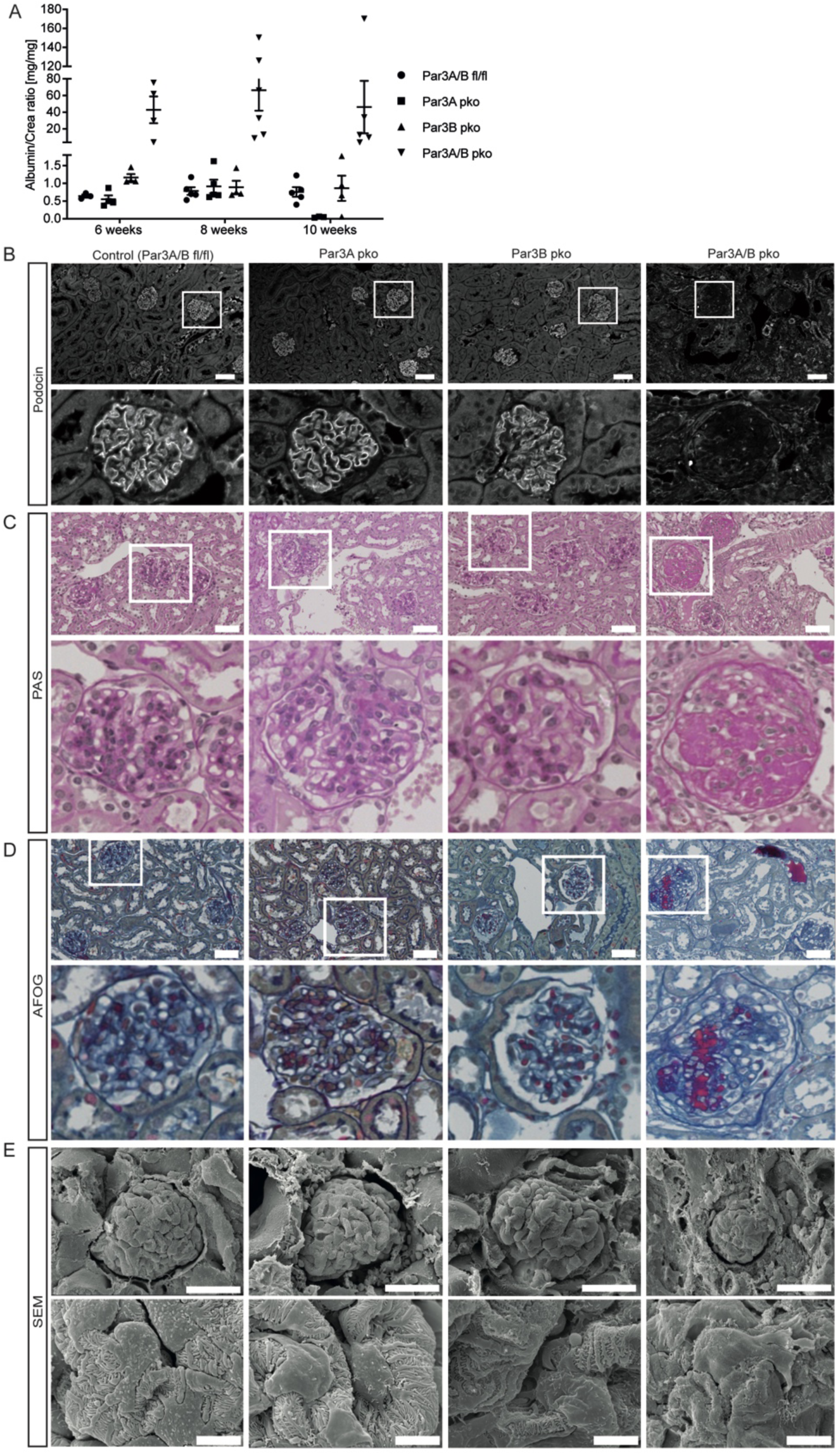
Loss of both, Par3A and Par3B, causes a severe glomerular disease phenotype. **A** Urinary albumin levels show an increase after 6 weeks only in Par3A/B double knockouts. Loss of Par3A or Par3B alone did not result in increased albumin levels. **B** Immunofluorescence stainings visualizing Podocin revealed complete disruption of the slit diaphragm upon loss of both, Par3A and Par3B.scale bar = 50**μ**m. **C** Histological analysis by performing periodic acid staining revealed severe glomerulosclerosis and protein casts in the tubular system in Par3A/B double knockouts. Scale bar = 50**μ**m. **D** Histological analysis by performing AFOG (acidic fuchsine orange G) staining revealed a large amount of protein deposits and thickening of the glomerular basement membrane upon loss of Par3A and B. Scale bar = 50μm. **E** Scanning electron microscopy revealed severe morphological changes including foot processes effacement after loss of Par3A and Par3B, while the single knockouts did not present with any morphological abnormalities. Upper panel: scale bar = 30**μ**m, lower panel: scale bar = 5**μ**m.

### Loss of Baz causes severe morphological and functional disturbances in *Drosophila* nephrocytes

In order to characterize the cellular and molecular function of the Par complex in the podocyte we utilized the *Drosophila* nephrocyte model taking advantage that the fly has only one Par3 orthologue (*Baz*). Nephrocytes are morphological and functional orthologues of mammalian podocytes and are widely used to study evolutionary conserved mechanisms at the kidney filtration barrier ^15^. Bazooka (Baz) is expressed in nephrocytes from the late embryonic stage onwards, where it co-localizes with the Neph homolog Dumbfounded (Duf) and the Nephrin homolog Sticks-and-stones (Sns) (garland cells, 3^rd^ instar larvae) (Figure 2A,B). Baz localization is not restricted to the nephrocyte diaphragm, but also expands into the cell body along the lacunae membranes (Figure 2A,B). In order to determine the function of Baz we generated a nephrocyte-specific knock-down of Baz using RNAi and assessed the phenotype of the nephrocyte diaphragm by observing the localization of the diaphragm proteins Duf and Polychaetoid (Pyd, the *Drosophila* ZO-1 ortholog) with high-resolution microscopy. In control nephrocytes (Sns-GAL4/+) the nephrocyte diaphragm is visualized as a fingerprint-like pattern (similar to the SD pattern in podocytes, e.g. Supp. Figure 4D) (Figure 2C). Upon loss of Baz this fingerprint-like pattern was almost completely lost and the integrity of the nephrocyte diaphragm was severely disrupted (Figure 2C). Disruption to the nephrocyte diaphragm is confirmed by the severe mislocalization of Duf, which no longer localizes to the diaphragm at the periphery of the cell, and is instead found internally within the cell (Figure 2D). The nephrocyte foot processes in Baz depleted nephrocytes are also markedly reduced, which resembles podocyte foot process effacement in Par3A/B double knockout mice (Figure 2E). In line with the morphological changes, loss of Baz also caused a severe filtration/uptake phenotype (Figure 2F). Hence, loss of Baz resulted in a pronounced nephrocyte phenotype including morphological and functional disturbances. This phenotype is highly similar to that caused by loss of Par3A/B in mammalian podocytes.

**Figure 2:**
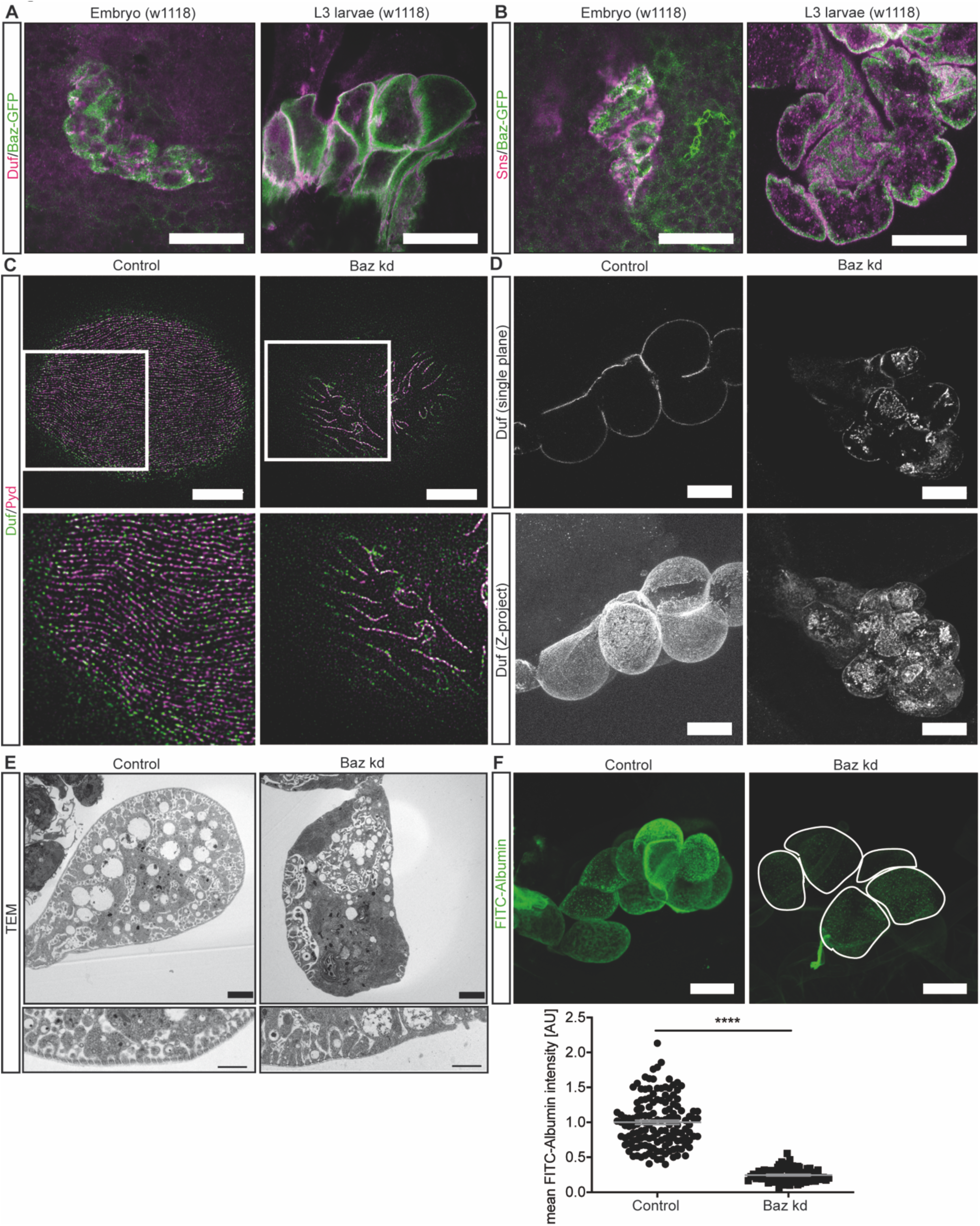
Loss of Baz causes severe morphological and functional disturbances in nephrocytes. **A,B** Immunofluorescence staining confirmed expression of Baz in wildtype nephrocytes (w1118), which are visualized by using a **A** Duf (dNeph) and a **B** Sns (dNephrin) antibody. Baz expression was assessed in garland nephrocytes of late stage embryos (first panel) and L3 larvae (second panel). Scale bar = 25**μ**m. **C** High-resolution microscopy revealed a finger-print like pattern in control cells (Sns-GAL4/+). Duf and Pyd (dZO-1) show an interdigitating pattern. Upon loss of Baz the finger-print like pattern is lost almost completely and the integrity of the nephrocyte diaphragm is completely disrupted. Scale bar = 5**μ**m. Control: w;Sns-GAL4;UAS-Dicer; Baz kd: w; Sns-GAL4/UAS-Baz-RNAi; UAS-Dicer/+. **D** Confocal microscopy shows an internalization of Duf upon loss of Baz (upper panel: single plane, lower panel: Z-projection) Scale bar = 25**μ**m. **E** Electron microscopy of control and Baz depleted nephrocytes shows a severe morphological defect in the Baz knockdown cells, including loss of foot processes and nephrocyte diaphragm structures. Scale bar = 2500nm. **F** FITC-Albumin filtration assays revealed a severe nephrocyte filtration defect after depletion of Baz. Students t-test: ****: p<0.005. Scale bar = 25**μ**m.

### Par3A and Par3B are both required for nephrocyte function

To investigate the compensatory effect of Par3A and B in greater detail, we tested the ability of different mammalian Par3 variants to rescue the *baz* knockdown phenotype (Supp. Figure 5). We investigated four different mammalian Par3A variants, of which three were detected in mouse podocytes, and the Par3B full length variant, which is the dominant variant in mouse podocytes (Figure 3A,B). Morphological assessment using high-resolution and electron microscopy, together with a functional analysis revealed different degrees of rescue potential (Figure 3C,D,E). We assessed the localization of Duf and Pyd by high-resolution microscopy and observed rescue with all three Par3A rescue strains and the Par3B rescue strain. The best rescue was found for the Par3A/B double rescue strain (Figure 3C). The Par3A 100kDa variant, which lacks the aPKC binding domain, also led to a partial rescue. Electron microscopy revealed a reduction of foot processes in Baz depleted nephrocytes, which was rescued by the expression of the different mammalian Par3 variants (Figure 3D). In addition, we performed functional analyses and investigated the filtration/uptake capacity of the different rescue strains (Figure 3E). Quantification revealed the best rescue in the Par3A/B double rescue strain (Figure 3E), superior to the full-length single rescue strains of Par3A and Par3B and the Par3A 100kDA variant (Figure 3E). In conclusion, the classical aPKC-binding domain of Par3 seems to be of minor importance for nephrocyte morphology and function as expression of the Par3 protein with no direct interaction with aPKC, i.e. Par3B, resulted in a comparable rescue to the Par3A full-length variant. This aligns with our finding that Par3B is sufficient for full function in podocytes.

**Figure 3:**
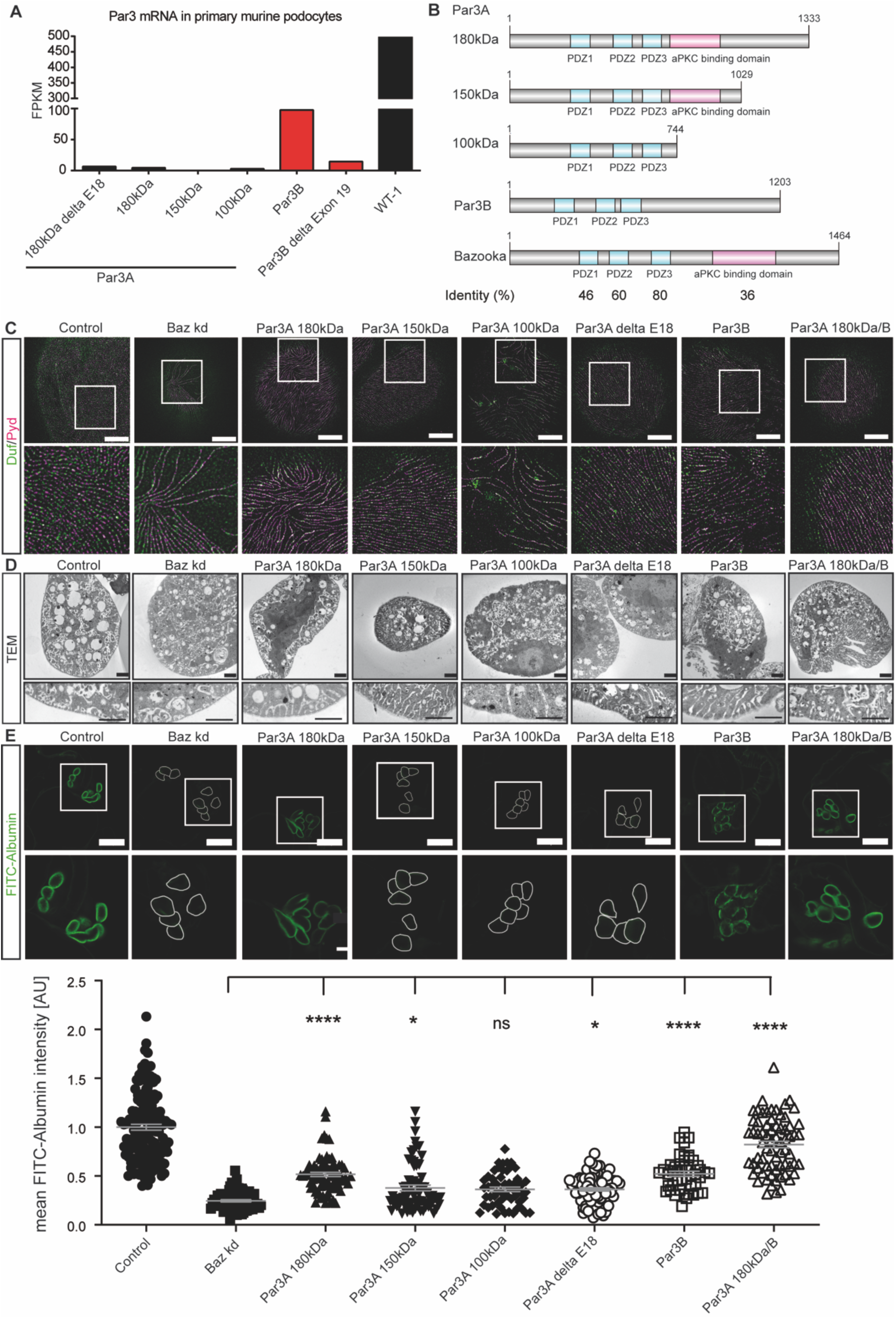
Par3A and Par3B are both required for proper nephrocyte function. **A** mRNA seq data from primary mouse podocytes revealed expression of three different Par3A variants (180kDa,100kDa and 180kDa delta Exon 18), while full-length Par3B was the most abundant Par3 protein ^53^. **B** Par3A, Par3B and Baz share high sequence similarity in the three PDZ domains. The Par3A 100kDa variant and Par3B do not have an aPKC binding domain. **C** High-resolution microscopy of all rescue fly strains shows the best rescue potential in the Par3A 180kDa/B double rescue fly strain. The nephrocyte diaphragm integrity was assessed by visualizing Duf and Pyd. The expression of the Par3A 100kDa variant, which lacks the aPKC binding domain, also resulted in a partial rescue, while expression of the two full-length variants resulted in a good and comparable rescue. Scale bar = 5μm. Control (Sns-GAL4/+), Par3A 180kDa (w;Sns-GAL4/UAS-Baz-RNAi;UAS-Par3A180kDa/UAS-Dicer), Par3A 150kDa (w;Sns-GAL4/UAS-Baz-RNAi;UAS-Par3A150kDa/UAS-Dicer), Par3A 100kDa (w;Sns-GAL4/UAS-Baz-RNAi;UAS-Par3A100kDa/UAS-Dicer), Par3A delta E18 (w;Sns-GAL4/UAS-Baz-RNAi;UAS-Par3A delta Exon 18/UAS-Dicer), Par3B (w;Sns-GAL4/UAS-Baz-RNAi;UAS-Par3B/UAS-Dicer), Par3A 180kDa/B (UAS-Par3B;Sns-GAL4/UAS-Baz-RNAi;UAS-Par3A180kDa/UAS-Dicer). **D** Performing electron microscopy confirmed the results from the high-resolution microscopy. The expression of both, Par3A and Par3B, resulted in the best rescue potential, when compared to Baz depleted nephrocytes. Scale bar = 2500nm. **E** To assess filtration function FITC-Albumin assays were performed and again showed, that the simultaneous expression of Par3A and Par3B resulted in the best rescue potential. Scale bar = 25μm. 1way ANOVA: *: p<0.01, ****: p< 0.005.

### Par3 function is mediated via aPKC-dependent and independent signaling pathways

As neither the Par3A 100kDa variant nor Par3B possess an aPKC binding domain but rescue the *baz* knockdown phenotype, it is suggested that Par3 is able to function independently from aPKC. aPKC is normally localized to the appropriate region in the membrane by binding the aPKC binding domain in Baz ^18–20^. We reasoned that analysis of an aPKC variant (aPKC-CAAX), that localizes independently of Baz, would allow us to uncouple the aPKC-dependent and independent functions of Baz. In addition, we also investigated the impact of the kinase domain on nephrocyte function by using a dominant negative aPKC-CAAX construct. The dominant negative construct harbors the K293W point mutation and leads to a non-functional kinase domain ^21^. To investigate the contribution of aPKC in the observed Baz phenotype, we first assessed nephrocyte morphology and second filtration function. With regard to morphology, expression of the aPKC-CAAX variant was able to restore Duf and Pyd localization comparable to the Par3A/B double rescue (Figure 4A). Expression of the dominant-negative variant resulted in a partial rescue, comparable to the Par3A 100kDA variant (Figure 3C, 4A). Transmission electron microscopy confirmed the rescue potential as seen in high-resolution microscopy (Figure 4B). However, functional analysis of the aPKC-CAAX rescue only revealed a partial rescue, which was significantly less efficient compared to the Par3A/B double rescue (Figure 4C). The aPKC-CAAX-DN variant showed the least rescue potential when compared to the Par3A/B double and aPKC-CAAX rescue (Figure 4C). Fly strains expressing either wildtype Baz or a mutant form lacking the aPKC-binding domain (Baz delta 968-996) in a *baz* knockdown background in nephrocytes were able to restore not only morphology as assessed by Duf and Pyd localization, but also filtration function (Figure 5A,B). Taken together, employing different aPKC and Baz mutants further emphasizes aPKC-independent functions of Par3A, Par3B and Baz.

**Figure 4:**
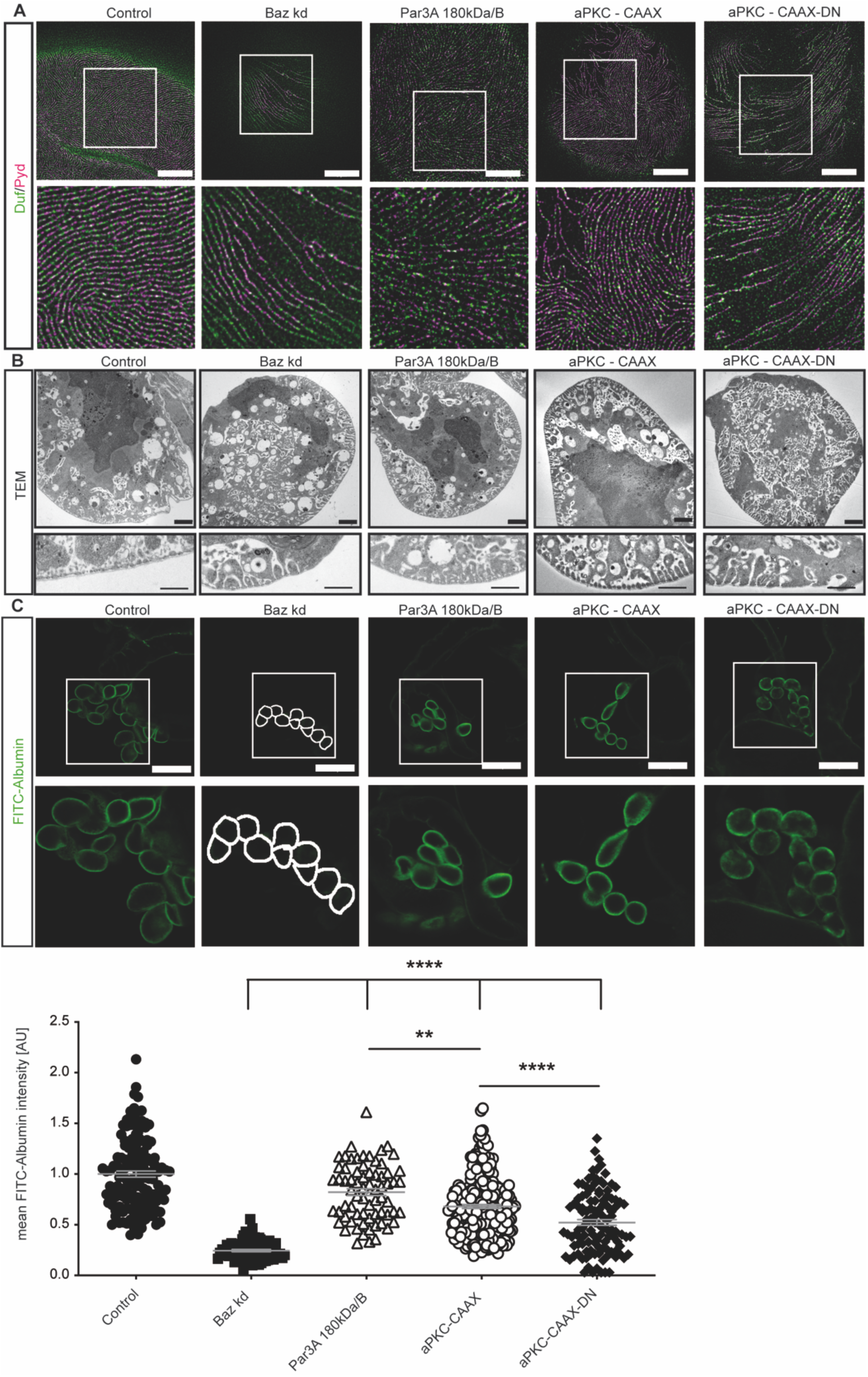
The kinase function of aPKC is required for proper nephrocyte morphology, but is less essential for filtration function. **A** High-resolution microscopy of nephrocytes expressing either aPKC-CAAX or aPKC-CAAX-DN in a Baz knockdown background shows a very good rescue capacity in the aPKC-CAAX nephrocytes, while expression of aPKC-CAAX-DN only resulted in a partial rescue. The resue capacity of aPKC-CAAX was comparable to the rescue of the Par3A/B double rescue fly strain. Scale bar =5μm. Control (w;Sns-GAL4/+;+), Baz kd (w;Sns-GAL4/UAS-baz-RNAi; Uas-Dicer/+), Par3A 180kDa/B (UAS-Par3B;Sns-GAL4/UAS-Baz-RNAi;UAS-Par3A180kDa/UAS-Dicer), aPKC-CAAX (w;Sns-GAL4/UAS-Baz-RNAi;UAS-aPKC-CAAX/UAS-Dicer), aPKC-CAAX-DN (w;Sns-GAL4/UAS-Baz-RNAi;UAS-aPKC-CAAX-DN/UAS-Dicer), **B** Electron microscopy confirmed the good rescue potential of aPKC-CAAX, while the expression of aPKC-CAAX-DN only caused a partial rescue. Scale bar = 2500nm. **C** FITC-Albumin filtration assays revealed a significant rescue in both, aPKC-CAAX and aPKC-CAAX-DN, fly strains, but also showed, that the double rescue with Par3A/B resulted in a significantly better rescue, when compared to the aPKC flies. Scale bar = 25μm. one-way-ANOVA: **: p > 0.01; ****: p > 0.001.

**Figure 5:**
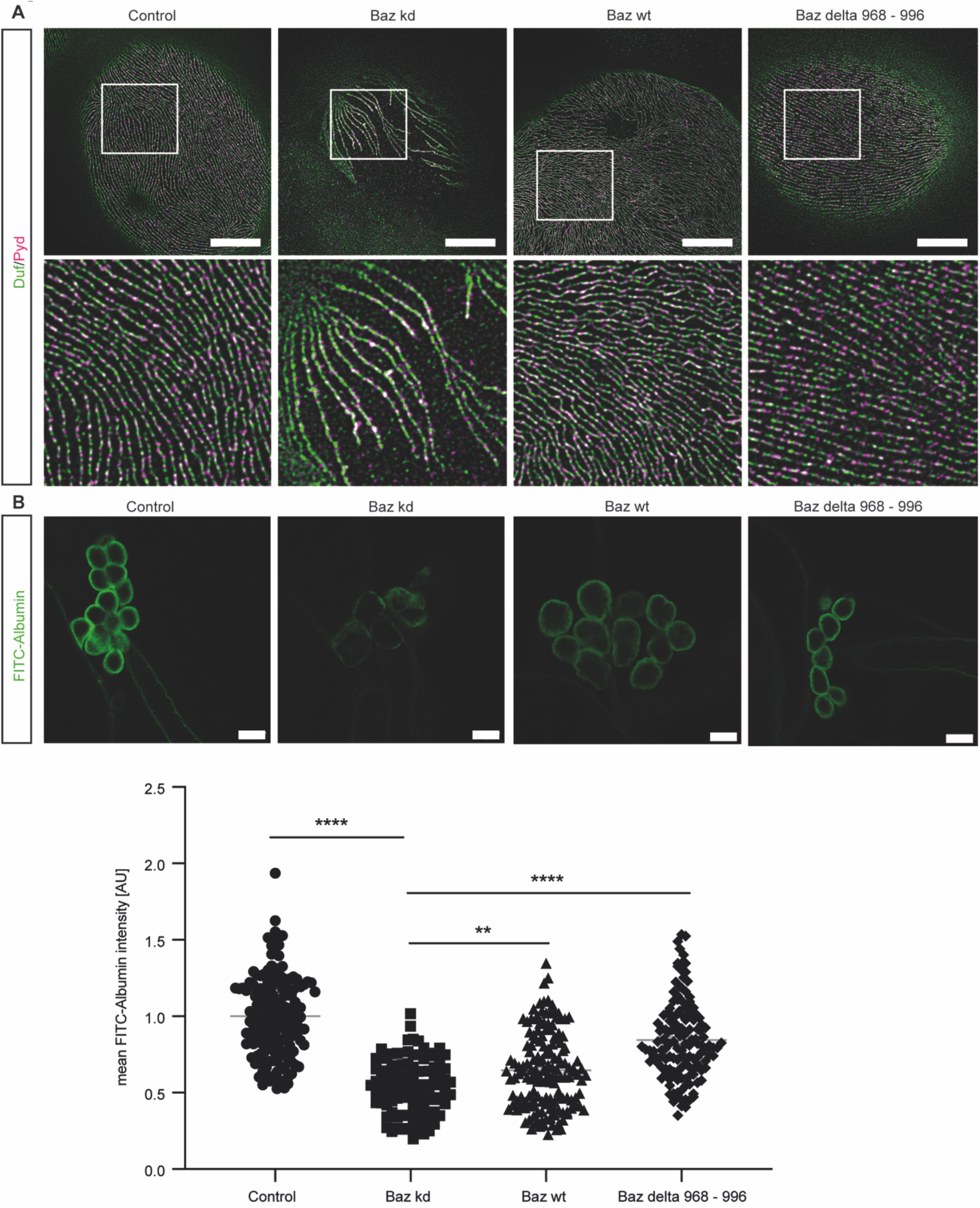
The aPKC-binding domain of Baz is dispensable for nephrocyte biology. **A** High-resolution microscopy using a Duf and a Pyd antibody revealed a very good rescue, when either wildtype Baz or a Baz mutant lacking the aPKC-binding domain, were expressed in a Baz knockdown background. Scale bar = 5μm. Control (w1118), Baz kd (Baz-GFP-trap; Sns-GAL4/+;UAS-GFP-RNAi/+), Baz wt (Baz-GFP-trap; Sns-GAL4/+;UAS-GFP-RNAi/UAS-Baz-wt), Baz delta 968 – 996 (Baz-GFP-trap; Sns-GAL4/+;UAS-GFP-RNAi/UAS-Baz-delta968-996). **B** FITC-Albumin filtration assays resulted in a partial, but significant rescue of both Baz expressing fly strains when compared to Baz depleted nephrocytes. The expression of the Baz delta 968-996 mutant caused an even better rescue in comparison to the wildtype Baz. Scale bar = 25μm. One-way ANOVA: **: p > 0.01; ****: p > 0.001.

### Tropomyosin2, the functional orthologue of Synaptopodin, is specifically up-regulated upon loss of Baz

Based on findings above we aimed to identify the aPKC-independent function of Baz in nephrocytes. In order to identify involved pathways downstream of Baz, we performed whole proteome analysis of isolated control and Baz depleted garland nephrocytes from 3^rd^ instar larvae and identified a significant upregulation of actin-cytoskeleton associated proteins (Actin, Tropomyosin), focal adhesion proteins (Vinculin) and matrix-associated proteins (Viking) (Figure 6A) which could be reversed by simultaneous expression of either Par3A or Par3B in the Baz knockdown background (Figure 6B,C,D). To investigate which of these candidates are specific to changes in Baz rather than more general changes associated with nephrocyte dysfunction we compared the Baz data set with whole proteome data sets from nephrocytes either depleted for Sns (Nephrin homolog) or Mec2 (Podocin homolog) (Supp Figure 6A,B,C). This comparison revealed a specific upregulation of these candidates upon loss of Baz and excluded a common upregulation observed in injured nephrocytes of different genetic backgrounds. We then focused on Tropomyosin2 (TM2) as functional orthologue of Synaptopodin, a critical actin regulator in podocytes ^22^. To provide further experimental evidence for the potential connection between TM2 and Baz, we generated interactome data sets of Par3A and Par3B in intramedullary collecting duct cells (IMCDs). Here, we identified unique and common interactors of the two Par3 proteins, among them CD2AP (Figure 6E,F, Supp Figure 7A,B), which was previously shown to interact with the TM2 orthologue Synaptopodin in podocytes ^23, 24^. Our data shows, a significant and specific upregulation of TM2 upon loss of Baz and provides evidence for a potential link of Par3/Baz and Synaptopodin/TM2 via CD2AP.

**Figure 6:**
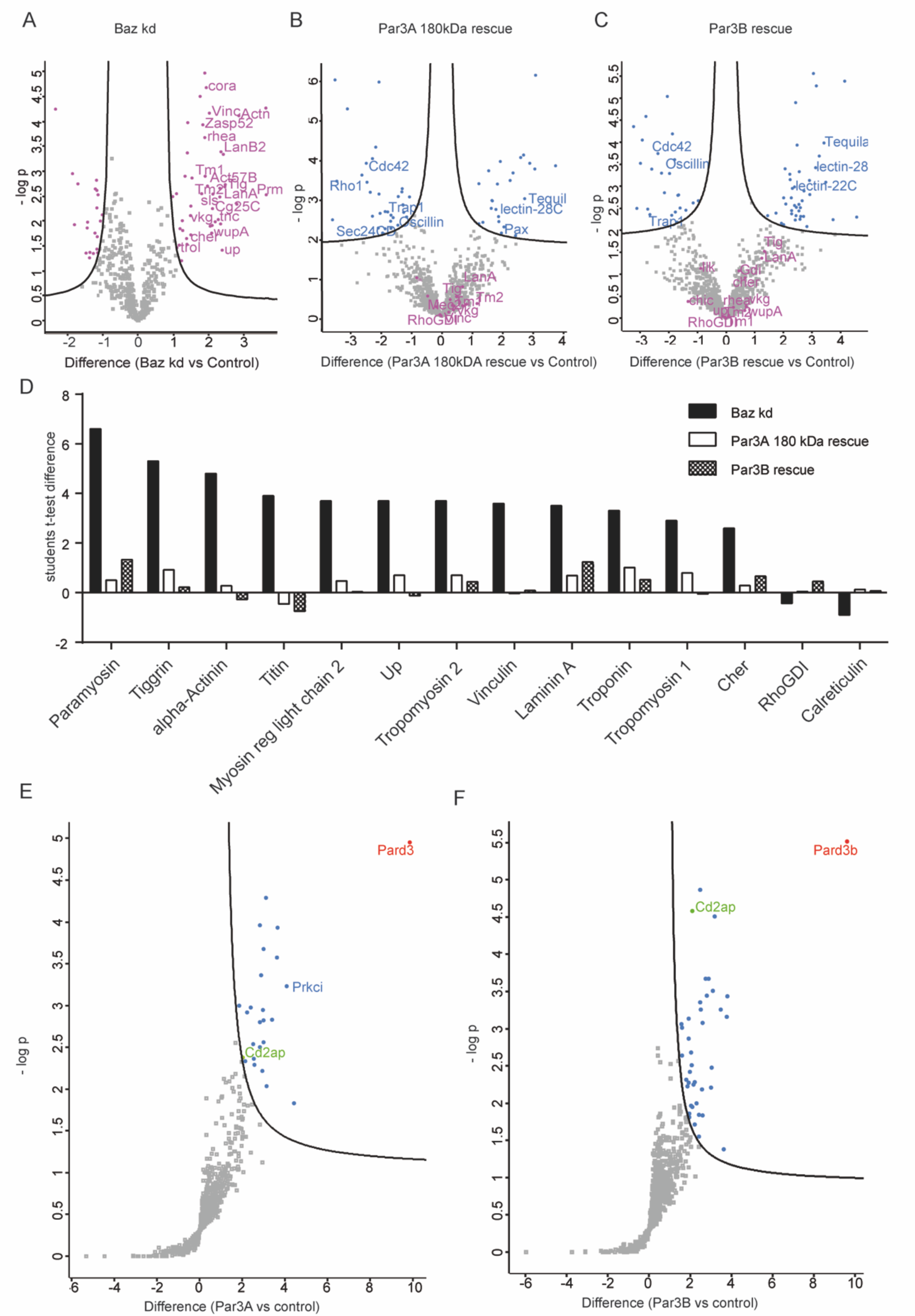
Tropomyosin2, the functional ortholog of Synaptopodin, is specifically up-regulated upon loss of Baz. **A** Volcano plot depicting differentially regulated proteins in Baz depleted nephrocytes compared to control cells. Isolated garland nephrocytes from single L3 larvae were used for proteome analysis. Significantly up-or down-regulated proteins are depicted in purple. FDR = 0.05, s0 = 0.1. **B,C** Volcano plot representing differentially regulated proteins in **B** Par3A 180kDa rescue nephrocytes and **C** Par3B rescue nephrocytes compared to control cells (Sns-GAL4/+). Blue: all proteins, which are significantly up- or down-regulated. Purple: proteins, which are not significantly regulated, but were identified as significantly regulated upon loss of Baz. FDR = 0.05, s0 = 0.1. **C** Comparison of interesting candidates of the Baz knockdown and the Par3A 180kDa and Par3B rescue proteome. Depicted are the students t-test differences. **D,E** Interactome analysis of **D** Par3A and **E** Par3B in intramedullary collecting duct cells. As control we used flag expressing cells. aPKC was only identified in the Par3A interactome, further confirming that there is no interaction between Par3B and aPKC. CD2AP (green) was identified as common interactor and potential link between Synaptopodin and Par3A and Par3B. Red: Par3A or Par3B, Blue: significant interactors, FDR = 0.05, s0 = 0.8.

### Baz regulates Rho1-GTP levels via Tropomyosin2

Captivated by the specific upregulation of TM2 in Baz depleted nephrocytes and the identification of CD2AP as common interactor of Par3A and Par3B, we hypothesized a functional interaction of Baz and TM2 via CD2AP and a potential contributing effect of the elevated TM2 levels to the Baz phenotype. Investigation of Baz and TM2 double knockdown fly strains, revealed a partial rescue regarding morphology and filtration function in comparison to the Baz single knockdown while loss of TM2 alone did not result in morphological disturbances and only showed a mild filtration defect when compared to Baz knockdown nephrocytes (Figure 7A,B). It was previously shown, that Synaptopodin and its *Drosophila* orthologue TM2 regulate RhoA signalling (in *Drosophila* Rho1) ^22, 25^, wherefore we employed Rho1 sensor flies (Rok-RBD-GFP) to analyze active Rho1 levels (Supp Figure 8A). In line with this, by comparing control and Baz depleted nephrocytes, we identified a significant increase of active Rho1 levels in Baz depleted cells (Figure 7C).

**Figure 7:**
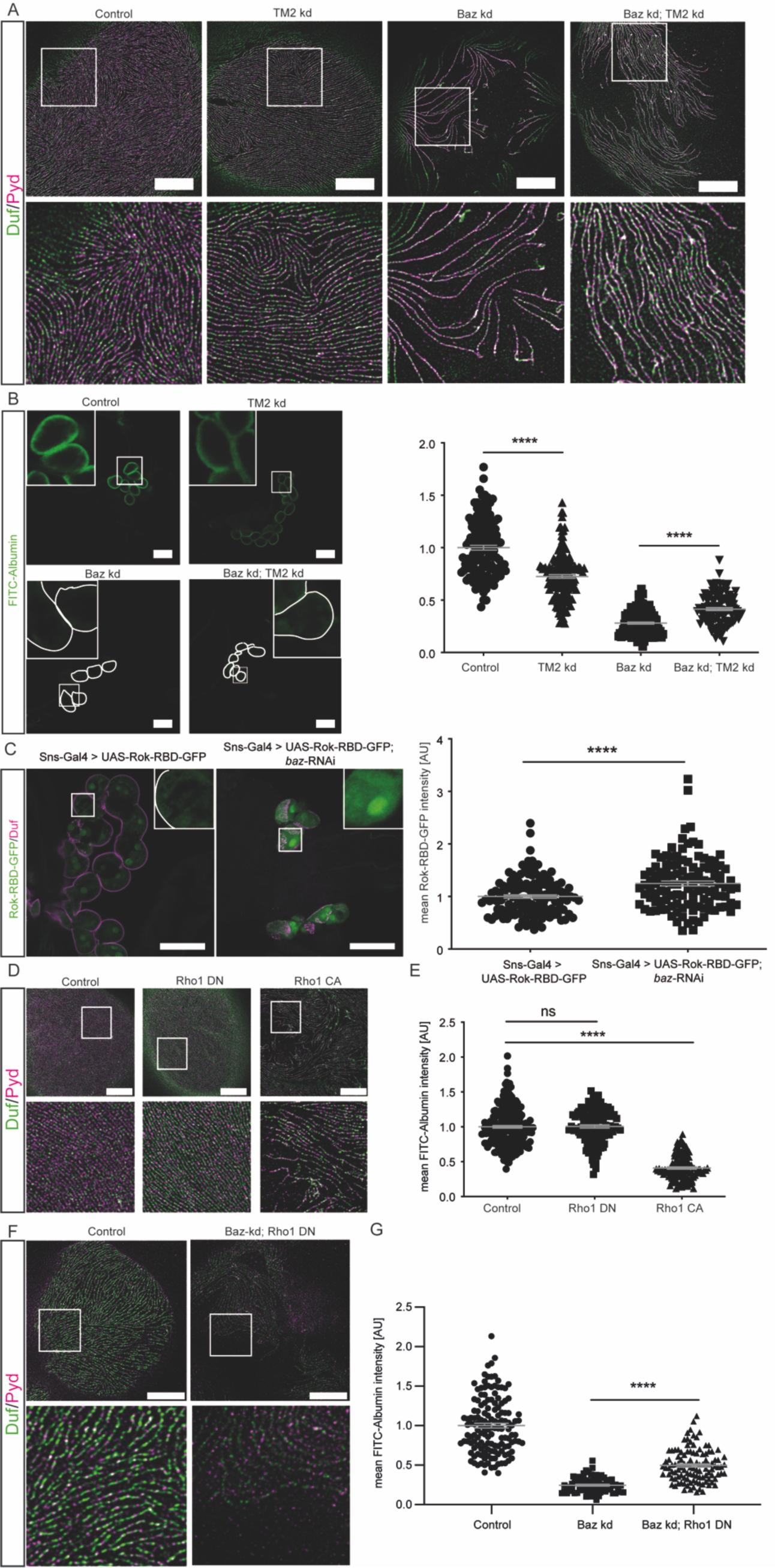
Baz regulates Rho1-GTP levels via Tropomyosin2. **A** High-resolution microscopy of control nephrocytes and nephrocytes depleted either for TM2, Baz or both, revealed a partial rescue upon simultaneous loss of both, Baz and TM2, in comparison to the Baz knockdown. Loss of TM2 alone did not result in a morphological phenotype. Scale bar = 5μm. Control (w;Sns-GAL4/+; UAS-Dicer/+), TM2 kd (w;Sns-GAL4/+;UAS-Dicer/UAS-*Tm2*-RNAi), Baz kd (w;Sns-GAL4/UAS-*baz*-RNAi; UAS-Dicer/+), Baz kd; TM2 kd (w;Sns-GAL4/UAS-*baz*-RNAi; UAS-Dicer/UAS-*Tm2*-RNAi). **B** FITC-Albumin assays confirmed the morphological phenotypes and showed a significant rescue in the Baz/TM2 double knockdown cells. Although loss of TM2 did not result in morphological abnormalities, it caused a significant reduction in filtration function. One-way ANOVA: ****: p < 0.001. **C** Confocal microscopy of Rho1 sensor flies showed a significant increase of GFP intensity in Baz depleted nephrocytes, proving elevated Rho1-GTP levels uponloss of Baz. Scale bar = 50μm. students t-test: ****: p < 0.001. **D** Expression of dominant negative Rho1 did not result in a morphological phenotype, while expression of constitutively active Rho1 caused severe morphological abnormalities as depicted by high-resolution microscopy. Scale bar = 5μm. Control: w;Sns-GAL4/+;UAS-Dicer/+, Rho1 DN: w;Sns-GAL4/+;UAS-Dicer/UAS-Rho1-DN, Rho1 CA: w;Sns-GAL4/+;UAS-Dicer/UAS-Rho1-CA. **E** FITC-Albumin filtration assays showed a severe filtration defect upon expression of constitutively active Rho1.Expression of dominant negative Rho1 did not result in any changes in comparison to the control nephrocytes. One-way ANOVA: ****: p > 0.001. **F** High-resolution microscopy revealed a partial rescue by expression of dominant negative Rho1 in a Baz knockdown background. Scale bar = 5μm. Control: w;Sns-GAL4/+;UAS-Dicer/+, Baz kd;Rho1 DN: w;Sns-GAL4/UAS-*baz*-RNAi;UAS-Dicer/UAS-Rho1-DN. **G** FITC-Albumin filtration assays revealed a significant and partial rescue of the filtration function upon expression of dominant negative Rho1 in a Baz knockdown background. One-way ANOVA: ****: p < 0.001.

To further assess the functional role of small GTPases in nephrocyte biology, we investigated the effects after expressing different GTPase variants in wildtype flies. Expression of a dominant-negative (DN) Rho1 variant did not result in a nephrocyte phenotype, while expression of a constitutively active (CA) Rho1 caused a severe morphological phenotype and filtration disturbances (Figure 7D). In addition, we also investigated the levels of active Rac1/Cdc42 upon loss of Baz, but did not observe any differences, when compared to control cells (Supp Figure 8B). To further strengthen the hypothesis of a functional interaction between Baz, TM2 and Rho-GTP levels, we generated flies expressing dominant negative Rho1 and *baz*-RNAi simultaneously. Intriguingly, expression of the dominant negative Rho1 variant was able to restore morphological features and filtration function of the *baz* knockdown phenotype partially (Figure 7E). Taken together, we observed an elevation of Rho1-GTP levels after depleting Baz. In line with this, diminishing Rho1-GTP levels in *baz* knockdown nephrocytes results in a partial rescue of the phenotype.

## Discussion

In this study, we link elements of polarity signaling to actin regulators to support the cellular architecture of glomerular podocytes in order to maintain the kidney filtration barrier. We describe novel and redundant functions of Par3A and Par3B. Although Par3B is expressed at much higher levels in podocytes, only simultaneous loss of both Par3 proteins results in a severe glomerular disease phenotype. This was recapitulated in the *Drosophila* nephrocyte model by depleting the Par3 homolog Baz. Mechanistically, both, Par3A and Par3B, interact with the actin regulator CD2AP independent of aPKC binding. Loss of the Par3 homolog Baz caused a significant increase of Synaptopodin/TM2 levels and elevated Rho-GTP levels resulting in podocyte and nephrocyte dysfunction.

The stringent control of GTPase signaling in podocytes in mice and humans is of central importance for podocyte function ^26–30^. Expression of both, dominant negative or constitutively active RhoA results in proteinuria and foot process effacement, while the expression of constitutively active Rac1 results in a spectrum of disease including phenotypes resembling FSGS or minimal change disease ^26, 29^. Inhibition of ROCK, which is downstream of RhoA, protects kidney function, while patients harboring mutations in ARHGDIA, a Rho GDP-dissociation inhibitor, suffer from steroid-resistant nephrotic syndrome ^27, 28^. Based on these findings, the activation of Rho1 upon loss of Baz might be causative for the phenotype observed. Interestingly, epidermal deletion of Par3A resulted in an inactivation of RhoA in keratinocytes ^31^. This opposite effect might be explained by the post-mitotic nature of podocytes, as the upstream regulation of RhoA via Par3A in keratinocytes seems to be important to maintain mitotic accuracy ^31^. Given that podocytes are very different in shape to keratinocytes, one may also envision that cycling of RhoA between active/inactive states might be important in regulating the cytoskeleton to help maintain the podocyte architecture.

Small GTPases also play a crucial role in maintaining the actin cytoskeleton, as the expression of constitutively active RhoA causes actin polymerization ^26^. Within this study, we identified the actin-cytoskeleton associated protein CD2AP as a common interactor of both Par3 proteins, thereby linking polarity proteins to Synaptopodin and the podocytes’ actin-based cytoskeleton (Figure 8) ^23, 24^. CD2AP is linking the slit diaphragm complex via Nephrin with the actin cytoskeleton ^32^ and was shown to be essential for podocyte biology, as loss of CD2AP resulted in renal failure and nephrotic syndrome already evident at four weeks after birth ^33^. Loss of Synaptopodin, however, did not lead to a glomerular disease phenotype, but caused higher susceptibility to injury models ^34^. Interestingly, CD2AP and Synaptopodin exhibit a functional interaction, as bigenic heterozygosity leads to proteinuria and a FSGS-like phenotype in mice ^23, 24^. The functional role of CD2AP is highly conserved throughout evolution as shown for Cindr the CD2AP homologue in *Drosophila* ^35^, further supporting our proposed mechanism including TM2/ Synaptopodin and Cindr/CD2AP. A current study investigated the impact of phosphorylation events of CD2AP and could show, that a tyrosine phosphorylation of CD2AP affects slit diaphragm stability ^36^. Whether phosphorylation events are also causative for the increased TM2/ Synaptopodin levels remain unclear for now.

**Figure 8:**
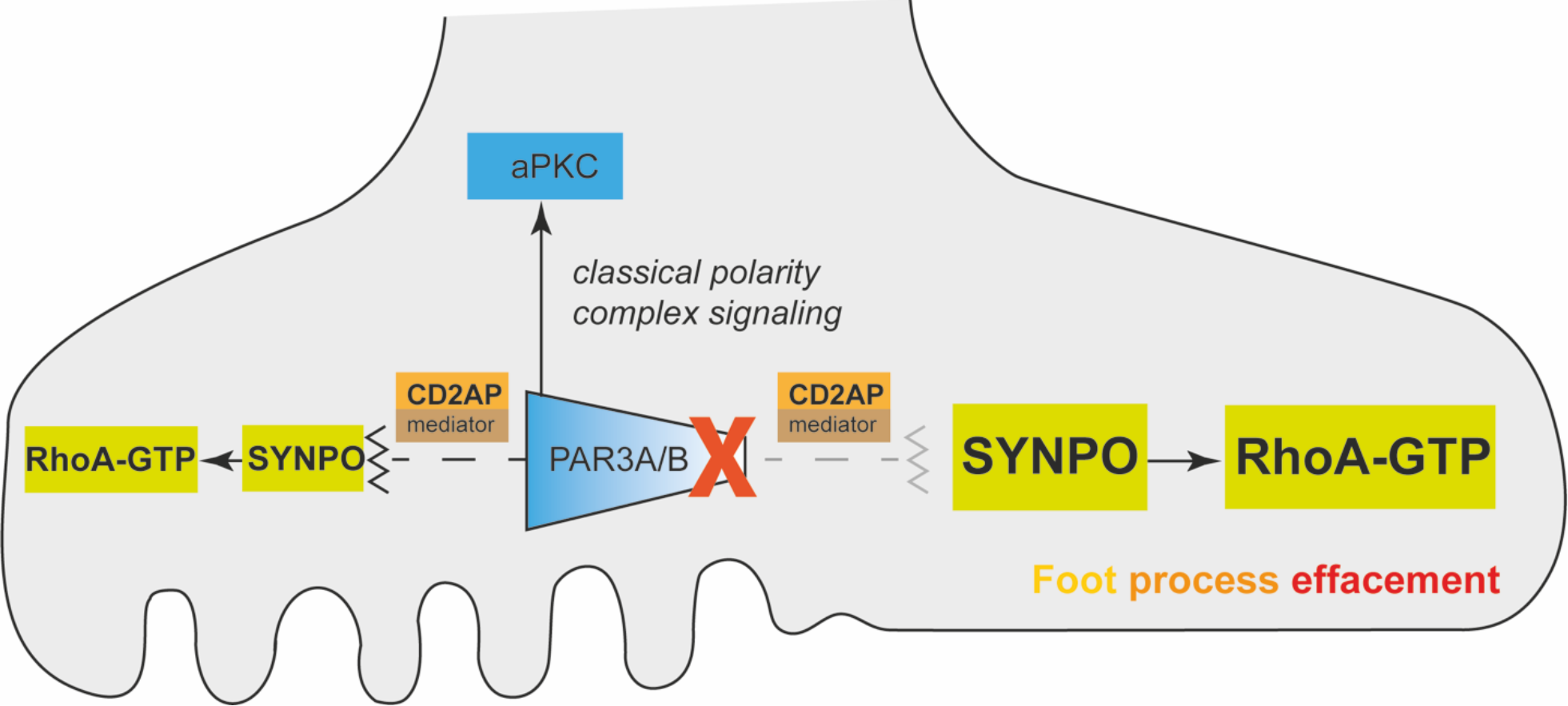
Par3 polarity proteins are linked to the actin cytoskeleton via CD2AP and are important to regulate Rho-GTP levels. The left half shows the healthy state, while the right half depicts the diseased condition after loss of Par3A/B in podocytes. The zig-zag line indicates that Par3A/B keep the mediator CD2AP under control.

Due to the podocytes’ unique morphology several studies have been performed to investigate the functional role of different polarity proteins belonging to the three major polarity complexes (Par, Crumbs and Scribble). Although loss of aPKCiota and Crumbs resulted in a disease phenotype in mice, zebrafish and *Drosophila* ^12, 37, 38^, depletion of Scribble and Par3A remained without any overt phenotype ^9, 10^. As podocytes only exhibit a single type of cell-to-cell contact combining properties of tight and adherens junctions, one can easily envision, that polarity complexes localize and function differently in comparison to classical epithelial cells. Further studies need to be done to unravel these potential redundant interactions of the different polarity proteins.

Our data emphasizes the importance of aPKC domains such as the PB1 domain for nephrocyte function, as the expression of a kinase-dead membrane-associated aPKC variant (aPKC-CAAX-DN) also resulted in a partial rescue of the *baz* knockdown phenotype. Kinase activity independent functions of aPKC have been described before in other *Drosophila* tissues, including the follicular epithelium ^39^. As aPKC has several protein-protein interaction domains, it may function as a scaffold to bring together proteins important for slit diaphragm formation.

The functional role of Par3B is only poorly understood. Studies in mammary gland stem cells revealed a unique interaction of Par3B with the tumor suppressor LKB1, which results in the inhibition of its kinase activity ^40, 41^. In contrast, Par3A does not seem to have a function in mammary gland stem cells ^40^. The highly similar protein structure suggests mutual functions of both Par3 proteins ^14^. Here, we identified the first common interactor of Par3A and B, which is not part of the well-known Par complex and provided first evidence of such a functional redundancy. In conclusion, Par3A and Par3B, are critical regulators of the podocyte architecture with high levels of redundancy. Mechanistically, Par3 proteins regulate Rho-GTP levels and interact with CD2AP independent of aPKC, thereby linking elements of polarity signaling to actin regulators to maintain the podocytes’ architecture.

## Material and Methods

### Antibodies

**Table 1:**
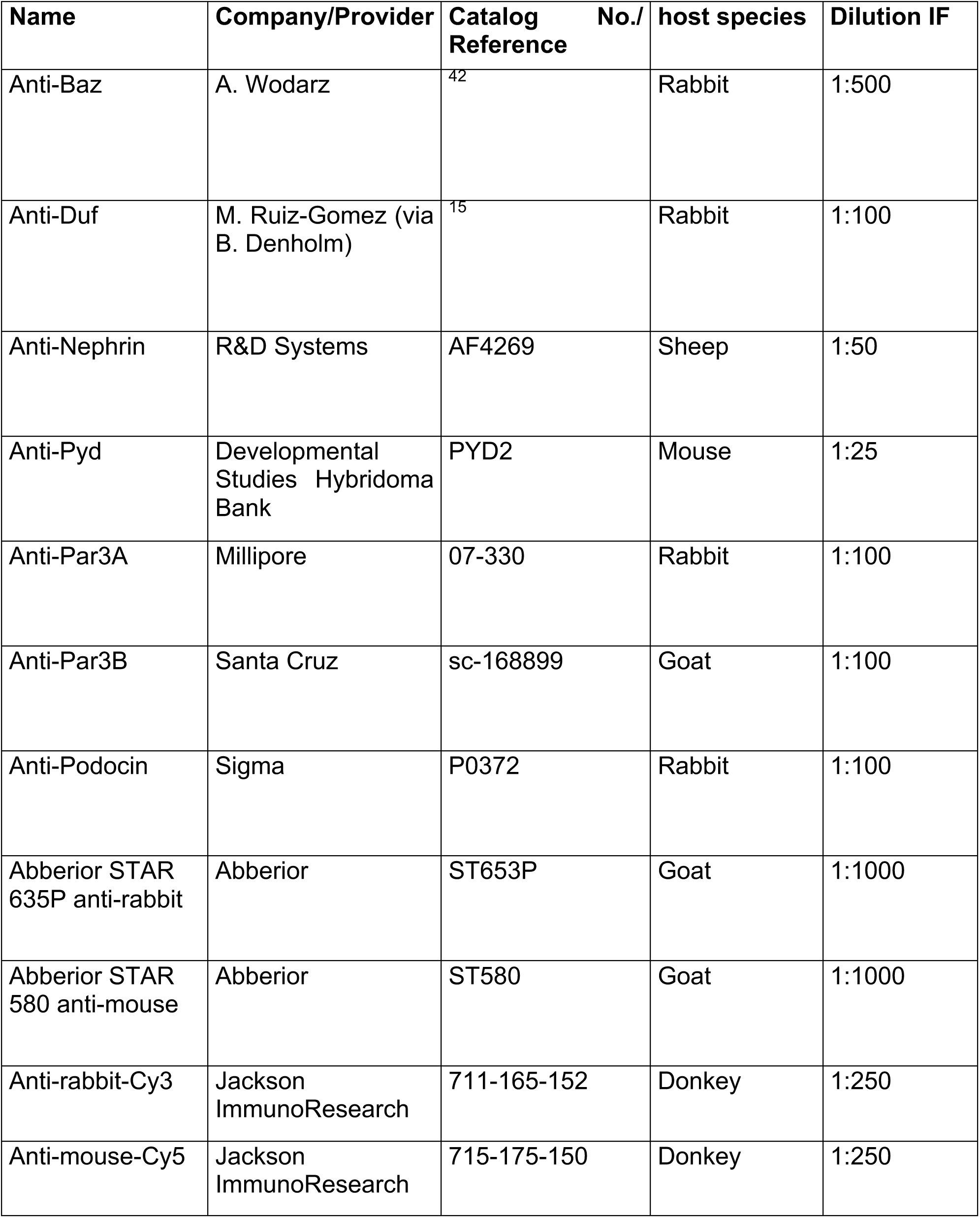
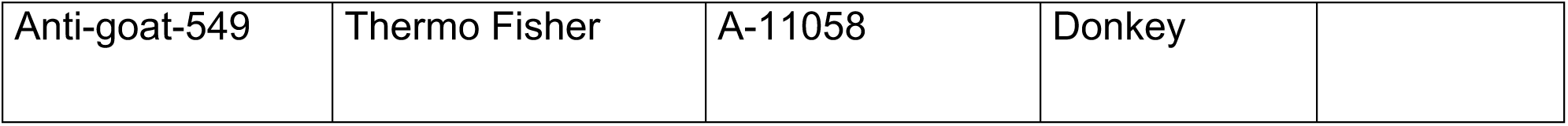
List of Antibodies.

### Primers

#### Deletion PCR Par3B

Par3B del for: TACCATTACCAGTTGGTCTGG

Pard3b-5’arm: GAAATGCCATATCCCTTTGTGACTGG

Pard3b-3’arm: CACAGCAACAGAACATGATTAATGGC

#### Quantitative PCR

mPar3B Sense: CGGTGCCATGCTGAGATTTG

mPar3B Anti-sense: TTCTGCTCAGCCTTTCCGAC

mACTB Sense: AAG AGC TAT GAG CTG CCT GA

mACTB Anti-sense: TAC GGA TGT CAA CGT CAC AC

#### Plasmids

Flag.Par3A pcDNA6.3 (selfmade)

Flag.Par3B pcDNA6.3 (selfmade)

Flag.pcDNA6.3 (selfmade)

### Cell culture experiments

For cell culture experiments, murine intramedullary collecting duct cells (IMCD) were used. They were cultivated with DMEM medium (Sigma-Aldrich, St. Louis, USA) with 5% FBS and 1% Glutamax. Cells were kept at 37°C with 5% CO2.

For transfection purposes cells were grown to a 60% density. Transfection was performed with GeneJuice (MerckMillipore, Darmstadt, Germany) according to manufacturer’s description using the plasmids listed in the plasmid section. After 48 h cells were lysed in immunoprecipitation (IP) buffer (20 mM Tris, 1% (v/v) Triton X-100, 50 mM NaCl, 15 mM Na4P2O7, 50 mM NaF, pH 7.5) with PIM (protease inhibitor mix) without EDTA (Roche, Basel, Switzerland). Lysates incubated with anti-flag M2 beads (Sigma-Aldrich, St. Louis, USA) for 1 h at 4°C. After washing with IP buffer, the beads were incubated with 80ul 5% SDS at 95°C for 5 min. The supernatant was reduced using 5 mM DTT for 30 min and alkylated using 10 mM IAA for 45 min at dark. Proteins were prepared and digested using Trypsin and LysC using the SP3 ultrasensitive proteomics technique as previously described ^43^.

### Fly stocks and husbandry

**Table 2:**
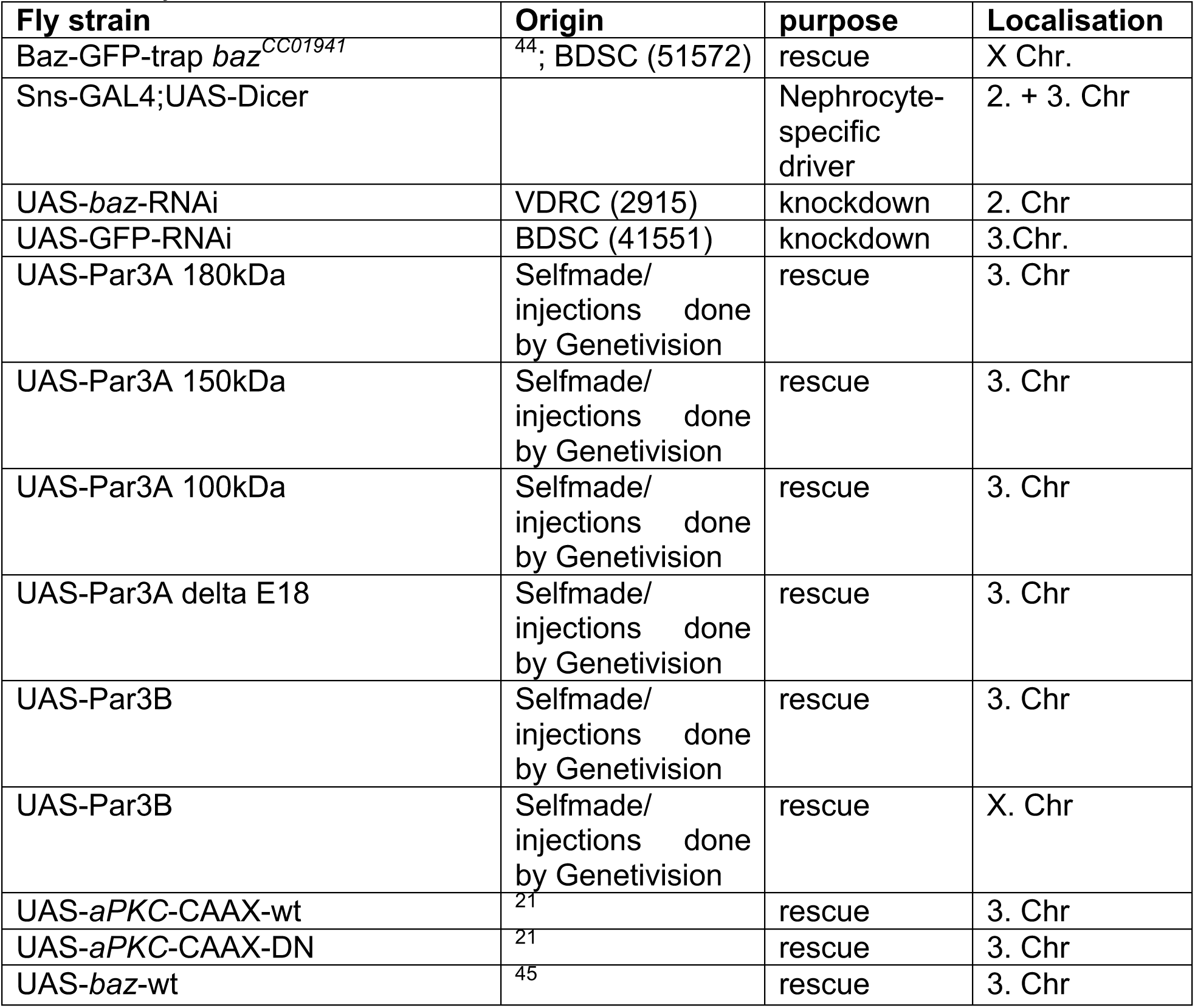
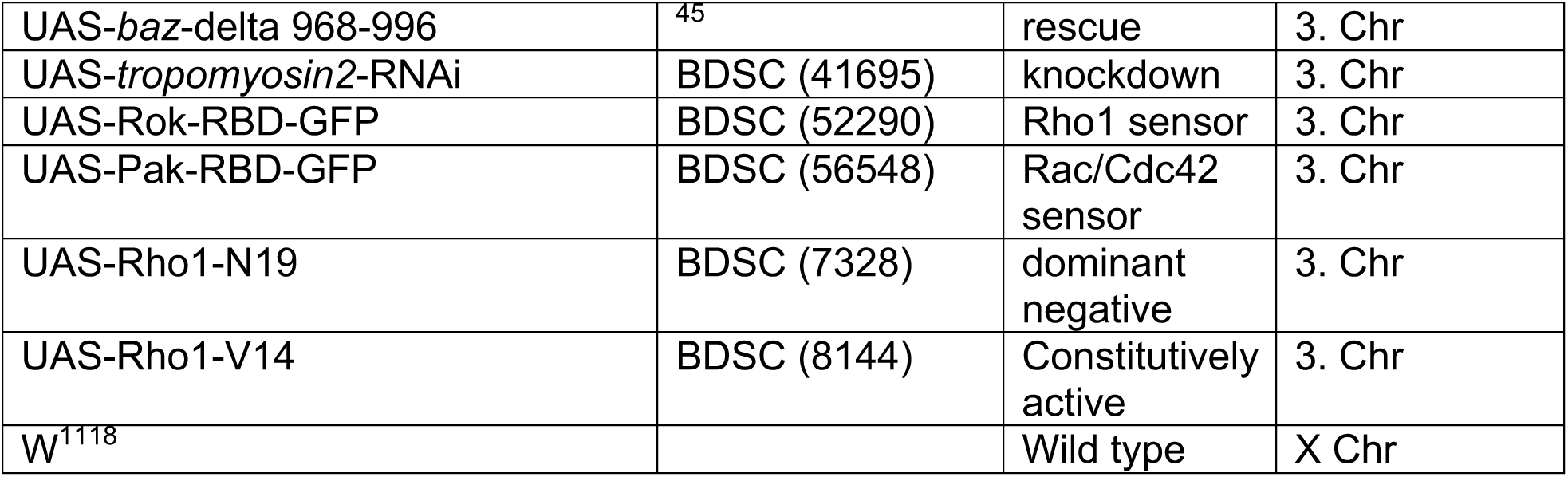
List of flystocks.

| **Fly strain** | **Origin** | **purpose** | **Localisation** |
| --- | --- | --- | --- |
| Baz-GFP-trap *baz*<sup>CC01941</sup> | <sup>44</sup>; BDSC (51572) | rescue | X Chr. |
| Sns-GAL4;UAS-Dicer |  | Nephrocyte-specific driver | 2. + 3. Chr |
| UAS-*baz*-RNAi | VDRC (2915) | knockdown | 2. Chr |
| UAS-GFP-RNAi | BDSC (41551) | knockdown | 3. Chr. |
| UAS-Par3A 180kDa | Selfmade/<br>injections done<br>by Genetivision | rescue | 3. Chr |
| UAS-Par3A 150kDa | Selfmade/<br>injections done<br>by Genetivision | rescue | 3. Chr |
| UAS-Par3A 100kDa | Selfmade/<br>injections done<br>by Genetivision | rescue | 3. Chr |
| UAS-Par3A delta E18 | Selfmade/<br>injections done<br>by Genetivision | rescue | 3. Chr |
| UAS-Par3B | Selfmade/<br>injections done<br>by Genetivision | rescue | 3. Chr |
| UAS-Par3B | Selfmade/<br>injections done<br>by Genetivision | rescue | X. Chr |
| UAS-*aPKC*-CAAX-wt | <sup>21</sup> | rescue | 3. Chr |
| UAS-*aPKC*-CAAX-DN | <sup>21</sup> | rescue | 3. Chr |
| UAS-*baz*-wt | <sup>45</sup> | rescue | 3. Chr |
| UAS-*baz*-delta 968-996 | <sup>45</sup> | rescue | 3. Chr |
| UAS-*tropomyosin2*-RNAi | BDSC (41695) | knockdown | 3. Chr |
| UAS-Rok-RBD-GFP | BDSC (52290) | Rho1 sensor | 3. Chr |
| UAS-Pak-RBD-GFP | BDSC (56548) | Rac/Cdc42 sensor | 3. Chr |
| UAS-Rho1-N19 | BDSC (7328) | dominant negative | 3. Chr |
| UAS-Rho1-V14 | BDSC (8144) | Constitutively active | 3. Chr |
| W<sup>1118</sup> |  | Wild type | X Chr |

Flies were kept at 25°C for all experiments. To achieve a nephrocyte-specific knockdown, flies were mated with a Sns - (sticks and stones) GAL4-driver line.

### Transgenic mouse models

The Par3A^fl/fl^ mouse model was previously published ^9^ and kept in a pure CD-1 background. The Par3B^fl/fl^ mouse model was generated using embryonic stem cells from EUCOMM and crossed onto a pure CD-1 background. Double knockout animals were generated by mating and also kept in a pure CD-1 background. Mating with a podocyte-specific Cre mouse model ^46^ resulted in podocyte-specific knockouts of Par3A, Par3B and both together. For podocyte isolation floxed animals were mated with R26mTmG mice ^47^, which were mated with hNphs2.PodCre mice to achieve GFP expression exclusively in podocytes ^46, 47^. For all animal experiments shown, we used both genders. The mouse husbandry was done in the University of Cologne animal facility according to standardized specific pathogen-free conditions. The experimental protocol was approved by the LANUV NRW (Landesamt für Natur, Umwelt und Verbraucherschutz Nordrhein-Westfalen, State Agency for Nature, Environment and Consumer Protection North Rhine-Westphalia, AZ 84-02.04.2013.A375).

### Isolation of primary podocytes

Primary podocytes were isolated after sacrificing mice and glomerular preparation as previously described ^48^. Glomeruli were either digested to get a single cell suspension, which was further used for FACs sorting or were used for DNA isolation using the Blood and Tissue kit from Qiagen (Hilden, Germany) according to manufacturer’s instructions.

### Immunofluorescence stainings and high-resolution imaging of *Drosophila* nephrocytes

Isolated nephrocytes were fixed with 4 % paraformaldehyde for 20 min followed by fixation in methanol for 2 h at room temperature. Afterwards they were washed three times in washing buffer (phosphate-buffered saline with 0.5 % BSA and 0.3 % Triton-X). Antibodies were used as described in table 1 and incubated overnight at 4°C. On the second day, nephrocytes were washed three times, blocked with 5 % normal donkey serum for 30 min and incubated with secondary antibody (see table 1, either for normal confocal or superresolution microscopy) for 1 h at room temperature and in the dark. Afterwards nephrocytes were washed three times and mounted using vectashield. Confocal images were obtained using a Zeiss Confocal microscope LSM710/Axiobserver Z1 and further processed using ImageJ/Fiji 1.50f8 and Adobe Photoshop Version 11.0. Superresolution images were acquired using a STED microscope (TCS SP8 gSTED 3x, Leica Microsystems) equipped with a white light laser for excitation and sensitive hybrid detectors (HyDs) for time gated detection. A 100x oil immersion objective with a numerical aperture of 1.4 (PL Apo 100x/1.4 Oil STED, Leica Microsystems) was used and the acquired images were processed using the deconvolution software Huygens Essential (Scientific Volume Imaging, SVI).

### Electron microscopy of *Drosophila* nephrocytes

For structural analysis, garland cells were prepared from L3 larvae and fixed in 2.5% glutaraldehyde in HL3 buffer (70mM NaCl, 4mM MgCl_2_x6H_2_O, 115mM Sucrose, 5mM KCl, 10mM NaKO_3_, 5mM HEPES, pH7.1). After washing in 100mM phosphate-buffer (NaH_2_PO_4_, NaH_2_PO_4_, pH 7.2), the cells were post fixed in 2% osmium tetroxide in phosphate-buffer for 1 hr on ice. After several washings in phosphate-buffer and dH_2_O the specimens were dehydrated in increasing ethanol and embedded in Araldit using aceton as an intermediate solvent. Thin sections were stained with 2% uranyl acetate and lead citrate. 50-100 nm sections were observed under an EM 109 (Zeiss) transmission electron microscope at 80 KV. Further processing was done by using ImageJ/Fiji 1.50f8 and Adobe Photoshop Version 11.0

### Histological analysis on mouse tissue

For histology mice were sacrificed via perfusion with phosphate-buffered saline (PBS) via the heart. Kidneys were either fixed in 4 % paraformaldehyde overnight followed by embedding in paraffin or fixed in 4 % paraformaldehyde for 2 h and further kept in PBS at 4°C (for high-resolution imaging). Periodic acid staining (PAS) and trichrome (AFOG) stainings were performed on 2 µm thick sections according to standard methods. All images were taken with a Leica SCN400 slide scanner and further processed using Aperio ImageScope v12.0.1.5030.

For immunofluorescence stainings kidneys were embedded in OTC. 5 um thick sections were used for imaging. Tissue was fixed with 4% paraformaldehyde for 8 min followed by washing for three times with PBS + 1% Triton-X. Afterwards tissue was blocked with 5 % normal donkey serum for 30 min and subsequent primary antibody incubation was performed over night at 4°C (antibodies see table 1). After three washing steps the secondary antibody was incubated for 1 h at RT, followed by washing and mounting with Prolong DAPI (New England Biolabs, Ipswich, USA). Images were obtained using a Zeiss Confocal microscope LSM710/Axiobserver Z1. Images were further processed using ImageJ/Fiji 1.50f8 and Adobe Photoshop Version 11.0. Exposure time was identical for comparative analyses.

For high-resolution microscopy, fixed pieces of kidney were incubated at 4°C in hydrogel solution (HS) (4% v/v acrylamide, 0.25% w/v VA-044 initiator, PBS 1X) overnight. The gel was polymerized at 37°C for 3 h, and the presence of oxygen was minimized by filling tubes all the way to the top with HS. After that, kidney pieces were cut into 0.3 mm thick slices using a Vibratome. Slices were then incubated at 50°C in clearing solution (200 mM boric acid, 4% SDS, pH 8.5) for 24 h. Prior to immunolabelling, samples were washed in PBST (0.1% Triton-X in 1X PBS) for 10 min.

For all steps PBST (PBS 1X with 0.1% v/v Triton-X) was used as dilutant. Samples were incubated in primary antibody for 24 h at 37°C, and were then washed in PBST for 10 min at 37°C followed by secondary antibody incubation for 24 h at 37°C and washed for 10 min at 37°C prior to mounting. To stain for Nephrin, a sheep anti-Nephrin primary antibody and a donkey anti-goat alexa-594 conjugated secondary antibody was used. Samples were immersed in 80% (w/w) fructose with 0.5% (v/v) 1-Thioglycerol at 37°C for 1 h prior to imaging. Samples were then mounted in a glass bottom dish (MatTek P35G-1.5-14-C) and imaged using a Leica SP8 3X STED system. Slit diaphragm (SD) length per area was determined using ImageJ/Fiji. In brief, the fluorescent signal was enhanced using background subtraction, mean filtering and contrast enhancement. The SD signal was detected with the help of the ridge detection plugin similarly to previously published by Siegerist et al. ^49^.

### Electron microscopy of mouse tissue

Primary fixation was done with 2% glutaraldehyde and 4% formaldehyde in 0.1M phosphate buffer for 48 hrs. For transmission electron microscopy (TEM), the tissue was then post-fixed in 0,5% osmium tetroxide in ddH2O for 60 min on ice and then washed 6 times in ddH2O. The tissue was incubated in 1% aqueous uranyl acetate solution for 2 hours in dark and washed 2 times in ddH2O.

Dehydration was performed by 15 min incubation steps in 30%, 50%, 70%, 90% and 2x 100% ethyl alcohol (EtOH) and 2x 100% aceton. After embedding in Durcupan resin ultrathin sections were performed using a UC7 Ultramicrotome (Leica), collected on Formvar-coated copper grids. Post staining was done for 1 min with 3% Lead Citrate followed by imaging using a Zeiss Leo 912 transmission electron microscope.

For scanning electron microscopy (SEM), the fixated kidneys were dehydrated in 70%, 80%, 90% and 100% ethanol (each step for 1 h at room temperature) and incubated in a 1:1 solution of EtOH and hexamethyldisilazan (HMDS) for 30 min. After incubation in 100% HMDS the solvent was allowed to evaporate. The dehydrated tissue was mounted onto sample holders and sputtered with gold using a Polaron Cool Sputter Coater E 5100. The resulting samples were imaged using a scanning electron microscope (Leo 1450 VP scanning).

### Urinary analysis

Urinary albumin levels were measured with a mouse albumin ELISA kit (ICL/Dunn Labortechnik GmbH, Asbach, Germany), while urinary creatinine levels were measured with a urinary creatinine kit (Biomol, Hamburg, Germany). Urinary albumin levels were then normalized on the urinary creatinine.

### Deletion PCR with glomerular lysates

Glomerular DNA was used for PCR using the RedTaq Ready MasterMix (Sigma-Aldrich, St. Louis, USA) according to manufacturer’s instructions. Used primers are listed in the primers section. Cycling conditions were as followed: 95°C 3 min, 95°C 30 sec, 60°C 30 sec, 72°C 30 sec go to step 2, repeat 35 times, 72°C 3 min. DNA was seperated using agarose gel electrophoresis and UV light.

### RNA isolation and quantitative PCR of murine podocytes

Podocytes were lysed with 500ul Trizol (Sigma-Aldrich, St. Louis, USA) and RNA was isolated using the Direct-Zoll RNA MiniPrep Kit from Zymo (Irvine, USA) according to manufacturer’s instructions. Isolated RNA was used for cDNA synthesis by using the High Capacity cDNA RT-Kit (Applied Biosystems, Foster City, USA) according to manufacturer’s instruction. Quantitative PCR was performed using the SYBR Green Mastermix (Thermo Fisher, Waltham, USA) and primers specific for mPar3B and mACTB (listed in the primers section). Par3B mRNA level were normalized to mACTB.

### FITC-Albumin uptake assay

To analyze the uptake and filtration capacity of nephrocytes, L3 larvae were dissected and garland nephrocytes were isolated. They incubated in FITC-Albumin (0,2 mg/ml; Sigma-Aldrich, St. Louis, USA) for 1 min ^50^. Afterwards they were washed in HL3.1 buffer for 1 min and then fixed in 4 % formaldehyde for 20 min. Nephrocytes were mounted in vectashield (Vectorlabs, Burlingame, USA) Images were obtained with a Zeiss Confocal microscope LSM710/Axiobserver Z1 and further processed using ImageJ/Fiji 1.50f8 and Adobe Photoshop Version 11.0. Exposure time was identical for comparative analyses. The mean intensity of single nephrocytes was measured and the values of the control cells (Sns-GAL4) was used for normalization.

### Proteomic analysis of *Drosophila* nephrocytes

Garland nephrocytes from single larvae were isolated and directly transferred into 40 ul 8 % SDS buffer. Samples were further reduced with 5 mM DTT and alkylated with 10 mM IAA in the dark. Afterwards proteins were further prepared and digested with Trypsin and LysC applying the SP3 method for ultrasensitive proteomics. Proteomics acquisition was performed on a quadrupole Orbitrap hybrid mass spectrometer (QExactive Plus, Thermo) coupled to a easynLC as previously described using 1h gradients for nephrocytes ^51^.

### Statistical analysis

Statistical significance was evaluated using GraphPad Prism version 6 for Windows (GraphPad Software, San Diego, CA). All results are expressed as means ± SEM. For comparison of two groups students t-test was used. A *P*-value of < 0.05 was considered to be significant. For one independent variable 1Way ANOVA combined with Tukey’s multiple comparison test was performed. A *P*-value of < 0.05 was considered to be significant. For two independent variables 2Way ANOVA combined with Sidak’s multiple comparison test was applied and a *P*-value < 0.05 was considered significant.

## Author contribution

S.K. and P.T.B. conceived the study; S.K., J.O., D.U.J., C.J., F.G. and M.H. performed experiments; S.K., M.H., W.B. and P.T.B. analyzed the data; S.I. and C.N. contributed animal models; S.K. made the figures and drafted the paper; S.K., H.H.H., S.I., C.N., G.W., A.W., B.S., T.B., B.D. and P.T.B revised the paper; all authors approved the final version of the manuscript.

## Acknowledgement

We thank V. Ludwig for her outstanding contributions and technical support. The authors also thank A. Koeser, G. Rappl, the CECAD imaging, proteomics and *in vivo* facilities for their support. S.K. received funding from the German Research Foundation (KO 6045/1-1), the Else-Kröner-Fresenius Foundation (2017_A135), the Köln Fortune Program of the University of Cologne, Germany as well as the German Society of Nephrology (Fritz-Scheler-Stipend). This study was supported by the German Research Foundation: clinical research unit (KFO 329, BR 2955/8-1 to P.T.B, SCHE 1562/7-1 to B.S. and BE 2212/23-1 + 2212/24-1 to T.B.). P.T.B. was supported by a DFG fellowship (BR2955/6-1). The Anti-Pyd monoclonal antibody developed by Fanning ^52^ was obtained from the Developmental Studies Hybridoma Bank, created by the NICHD of the NIH and maintained at The University of Iowa, Department of Biology, Iowa City, IA 52242.

## Supplementary Figures

**Supplementary Figure 1:**
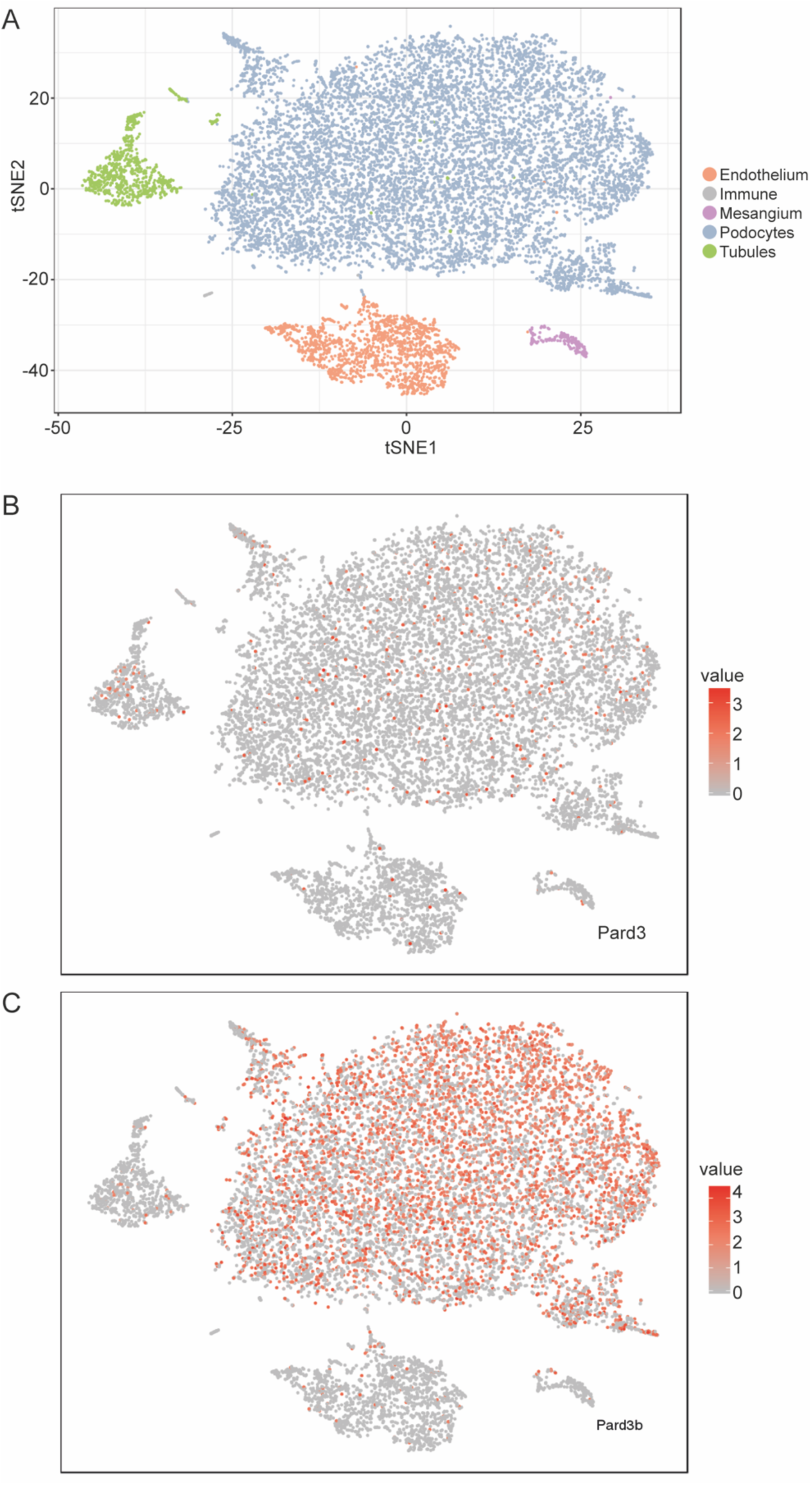
Single cell sequencing data from murine glomeruli. **A** Single cell sequencing of murine glomeruli revealed five subclusters of different cell types. **B** Pard3B was highly enriched in the podocyte subcluster, while **C** Pard3a was ubiquitously found.

**Supplementary Figure 2:**
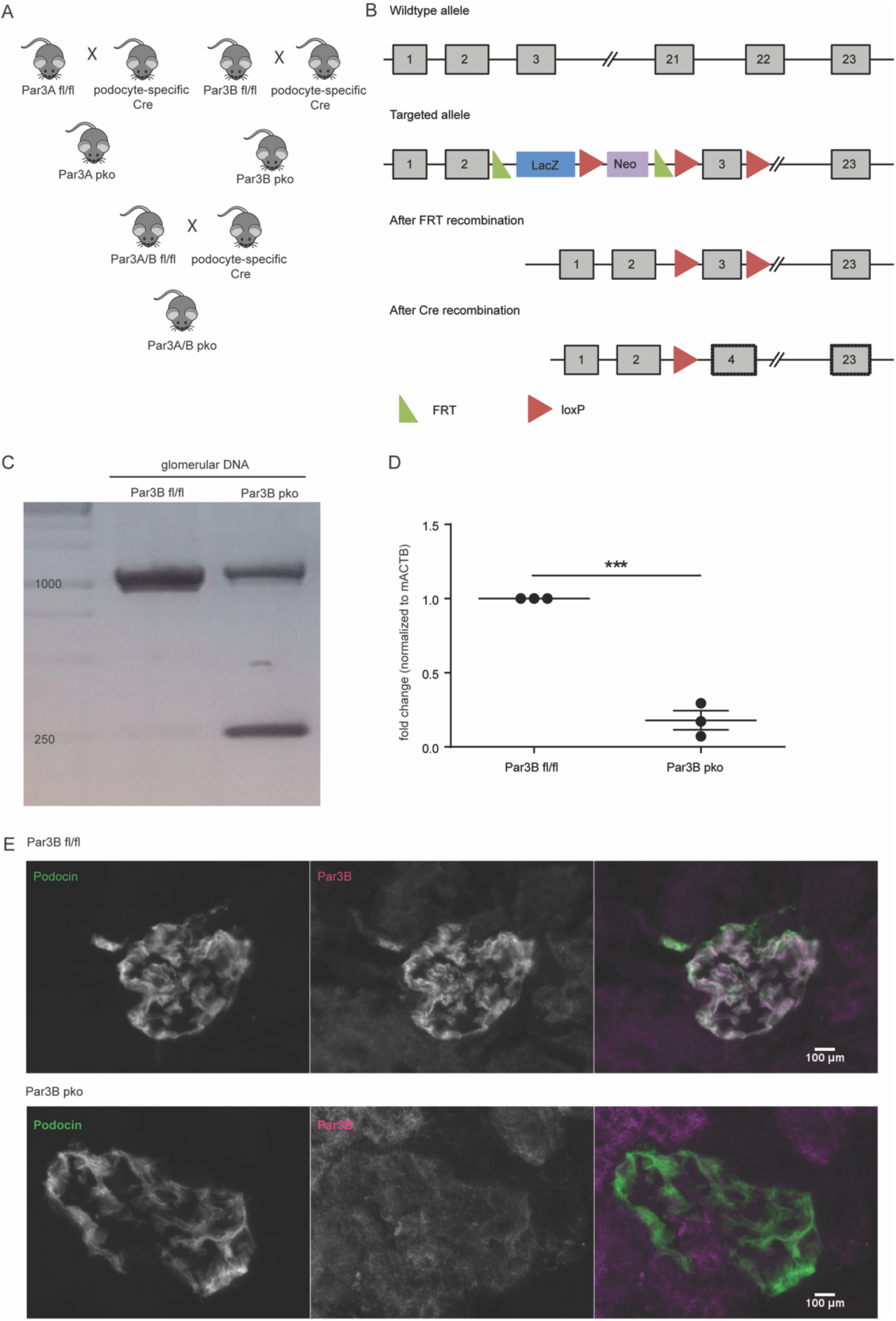
Generation of a novel podocyte-specific Par3B knockout mouse model. **A** Mating scheme to achieve podocyte-specific knockouts of Par3A, Par3B and Par3A/B double knockouts. **B** Scheme to present the generation and knockout strategy. Exon 3 was flanked by loxP sites, which results in a premature STOP after Cre-mediated excision. Embryonic stem cells were obtained from EUCOMM and used for *in vitro* fertilization. **C** Deletion PCR with glomerular DNA lysates revealed an additional band after Cre-mediated excision, which represents the truncated gene. As we used glomerular lysates other cell such as mesangial or endothelial cells still carry the wildtype gene, wherefore a wildtype band can be seen. **D** The remaining mRNA levels were investigated by quantitative PCR and showed a significant reduction of Par3B levels in primary mouse podocytes. For normalization ActinB primers were used. Students t-test: ***: p< 0.05. **E** Immunofluorescence stainings confirmed the loss of Par3B in podocytes. To visualize the slit diaphragm a Podocin antibody was used. Scale bar = 100**μ**m.

**Supplementary Figure 3:**
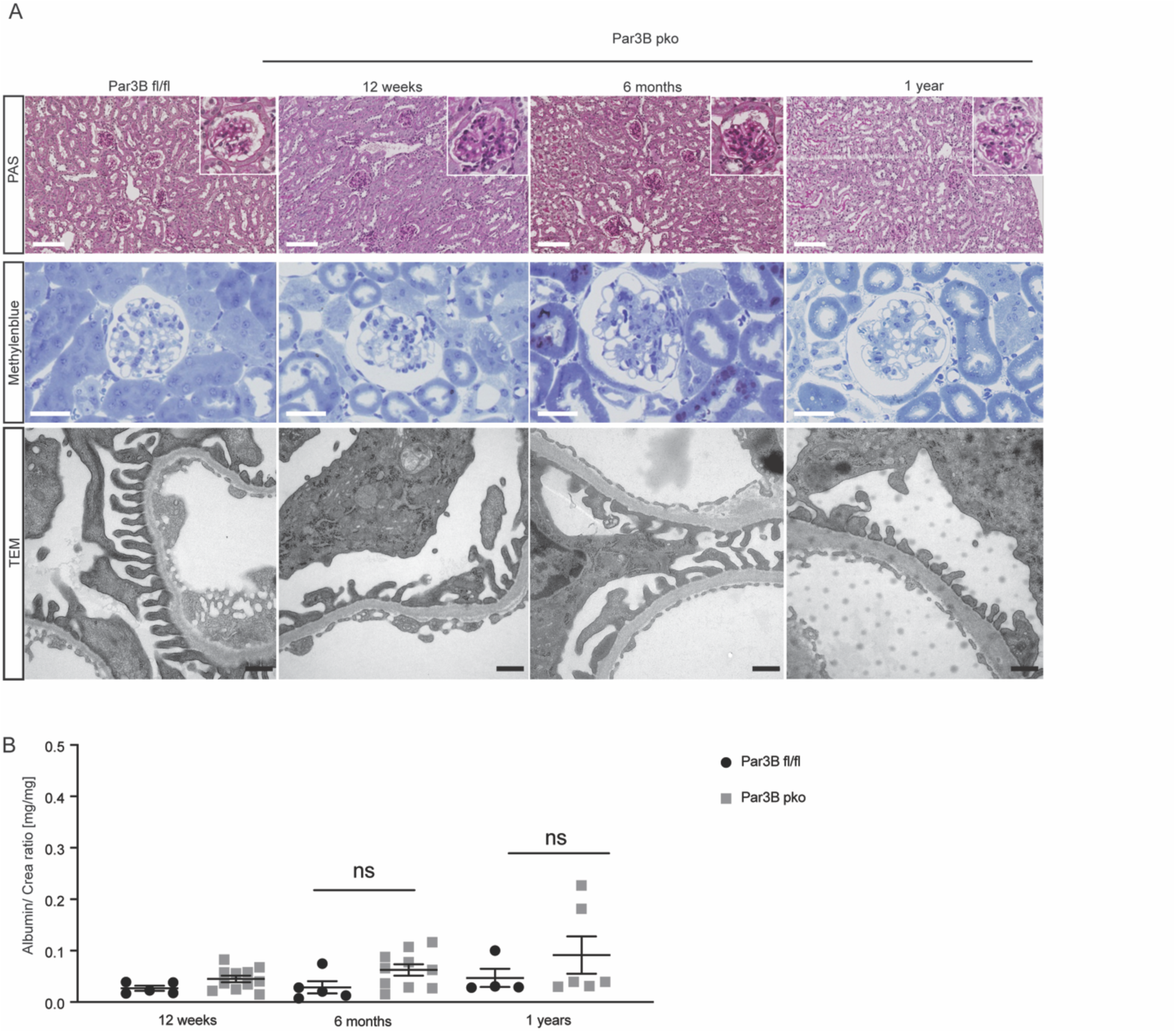
Loss of Par3B does not result in a glomerular disease phenotype. **A** Histological analysis such as periodic acid and methylene blue staining as well as electron microscopy did not reveal any differences between Par3B knockout mice and the control animals. Upper panel: scale bar =100**μ**m, middle panel: scale bar = 30**μ**m, lower panel: 1**μ**m. **B** Urinary albumin levels did not show any significant differences between podocyte-specific Par3B knockout and control animals for up to one year. One-way ANOVA with Sidak’s multiple comparison test.

**Supplementary Figure 4:**
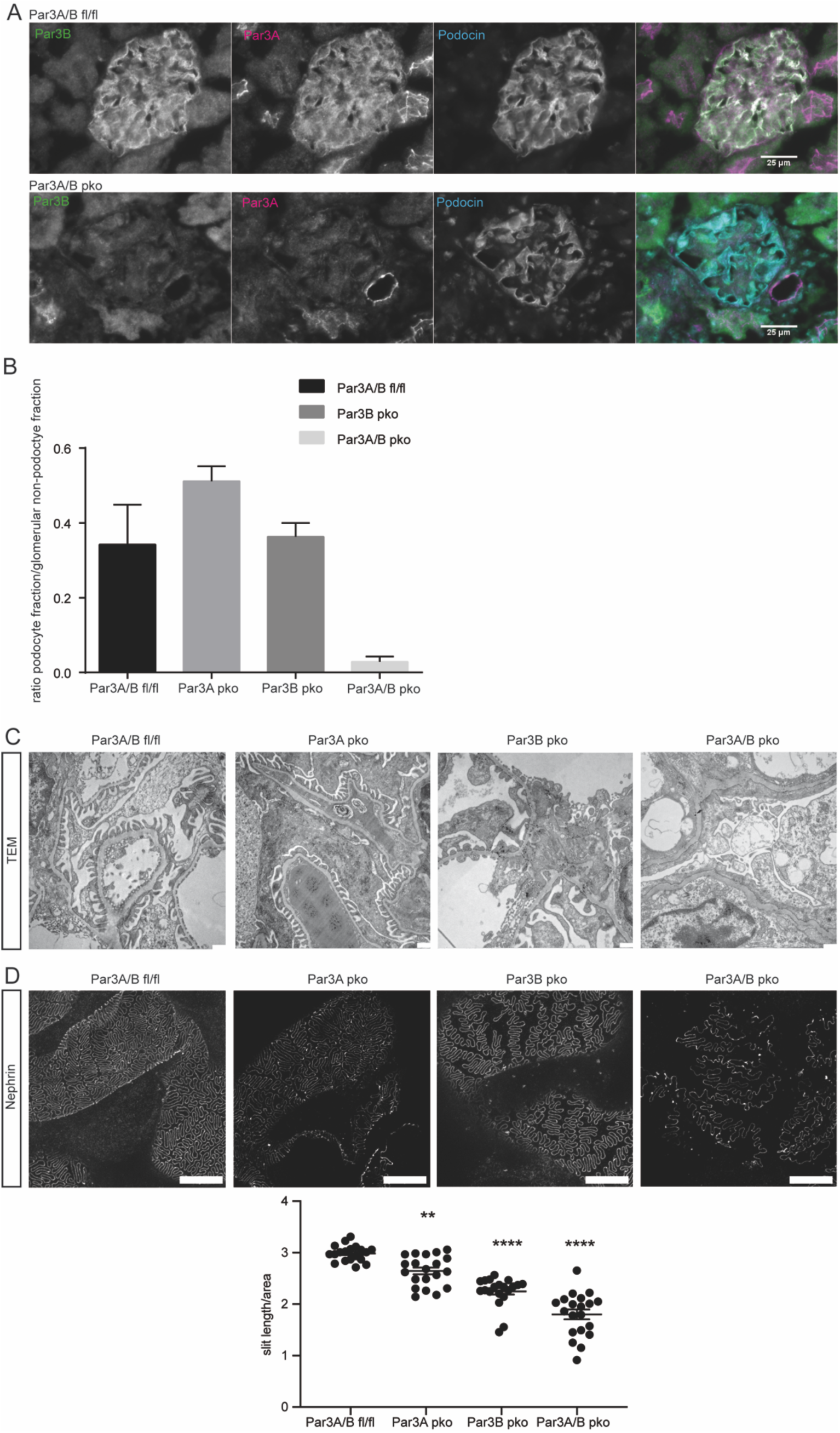
Generation of Par3A and Par3B double knockout mice. **A** Immunofluorescence staining revealed complete loss of Par3A and Par3B expression at the slit diaphragm, visualized by Podocin staining, in podocyte-specific Par3A/B double knockout mice. Scale bar = 25**μ**m. **B** FACS analysis of isolated primary podocytes revealed a severely decreased ratio of podocyte to non-podocyte fraction in Par3A/B double knockout cells. **C** Transmission electron microscopy revealed foot process effacement in Par3A/B double knockout mice, while depletion of Par3A and Par3B alone did not result in morphological abnormalities. Scale bar = 500nm. **D** High-resolution microscopy visualizing Nephrin showed a decreased slit diaphragm coverage in all three knockout models, while loss of both Par3 proteins caused the most severe phenotype. Scale bar = 5 um. 3 animals per genotype. One-way ANOVA with Tukey’s multiple comparison test: **: p < 0,005; ****: p < 0,0001.

**Supplementary Figure 5:**
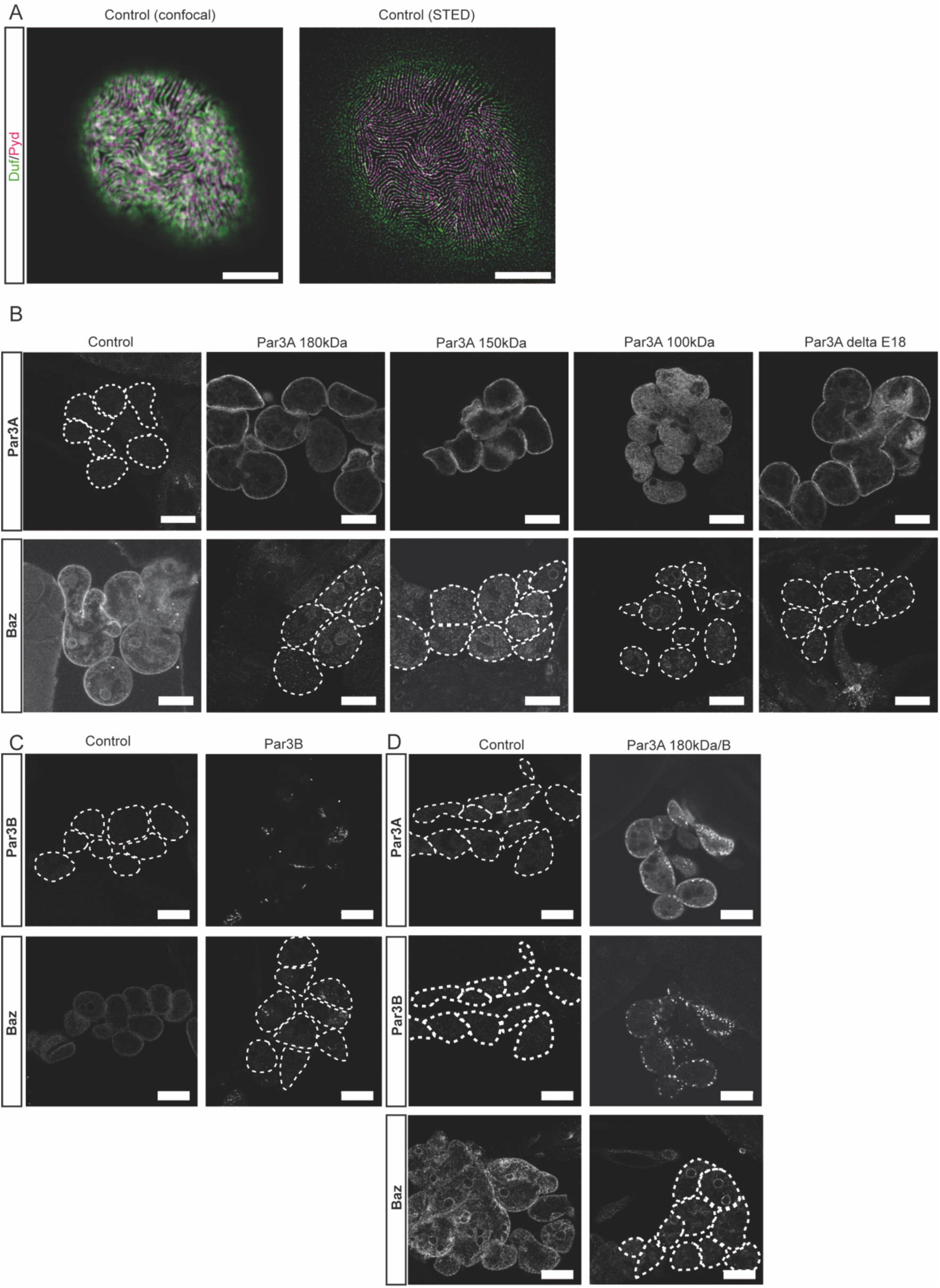
Generation of Par3 rescue fly strains. **A** Comparison of confocal and high-resolution microscopy. Scale bar = 5μm. **B** Four different Par3A variants are expressed in a Baz knockdown background. Expression of the different mammalian variants as well as the Baz knockdown was confirmed by immunofluorescence staining. Scale bar = 25μm. Control (Sns-GAL4/+), Par3A 180kDa (w;Sns-GAL4/UAS-Baz-RNAi;UAS-Par3A180kDa/UAS-Dicer), Par3A 150kDa (w;Sns-GAL4/UAS-Baz-RNAi;UAS-Par3A150kDa/UAS-Dicer), Par3A 100kDa (w;Sns-GAL4/UAS-Baz-RNAi;UAS-Par3A100kDa/UAS-Dicer), Par3A delta E18 (w;Sns-GAL4/UAS-Baz-RNAi;UAS-Par3A delta Exon 18/UAS-Dicer) **C** Expression of Par3B and simultaneous depletion of Baz was confirmed by immunofluorescence staining. Scale bar = 25μm. Control (Sns-GAL4/+), Par3B (w;Sns-GAL4/UAS-Baz-RNAi;UAS-Par3B/UAS-Dicer) **D** Par3A and Par3B expression as well as loss of Baz was confirmed by immunofluorescence staining. Scale bar = 25μm. Control (Sns-GAL4/+), Par3A 180kDa/B (UAS-Par3B;Sns-GAL4/UAS-Baz-RNAi;UAS-Par3A180kDa/UAS-Dicer).

**Supplementary Figure 6:**
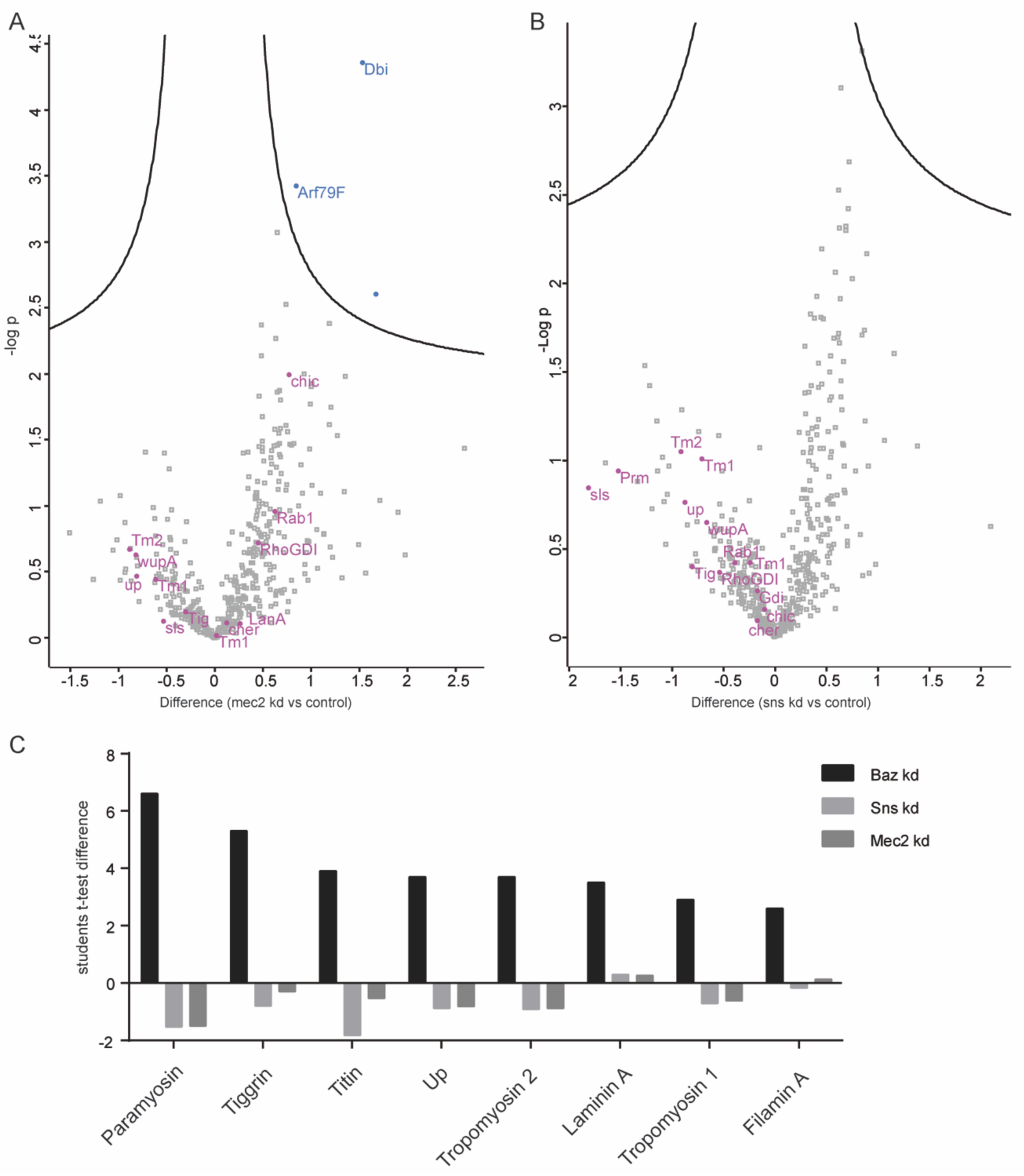
Whole proteome analysis of Sns and Mec2 depleted nephrocytes. **A** Volcano plot depicting differentially regulated proteins in Mec2 (dPodocin) depleted nephrocytes compared to controls (Sns-GAL4/+). Blue: significant proteins. Purple: proteins, which are significantly up-regulated in Baz knockdown nephrocytes. FDR = 0.05, s0 = 0.1. **B** Volcano plot depicting differentially regulated proteins in Sns (dNephrin) depleted nephrocytes compared to controls (Sns-GAL4/+). Purple: proteins, which are significantly up-regulated in Baz knockdown nephrocytes. FDR = 0.05, s0 = 0.1. **C** Bar graph comparing the students t-test difference of interesting candidates between Baz, Sns and Mec2 knockdown nephrocytes.

**Supplementary Figure 7:**
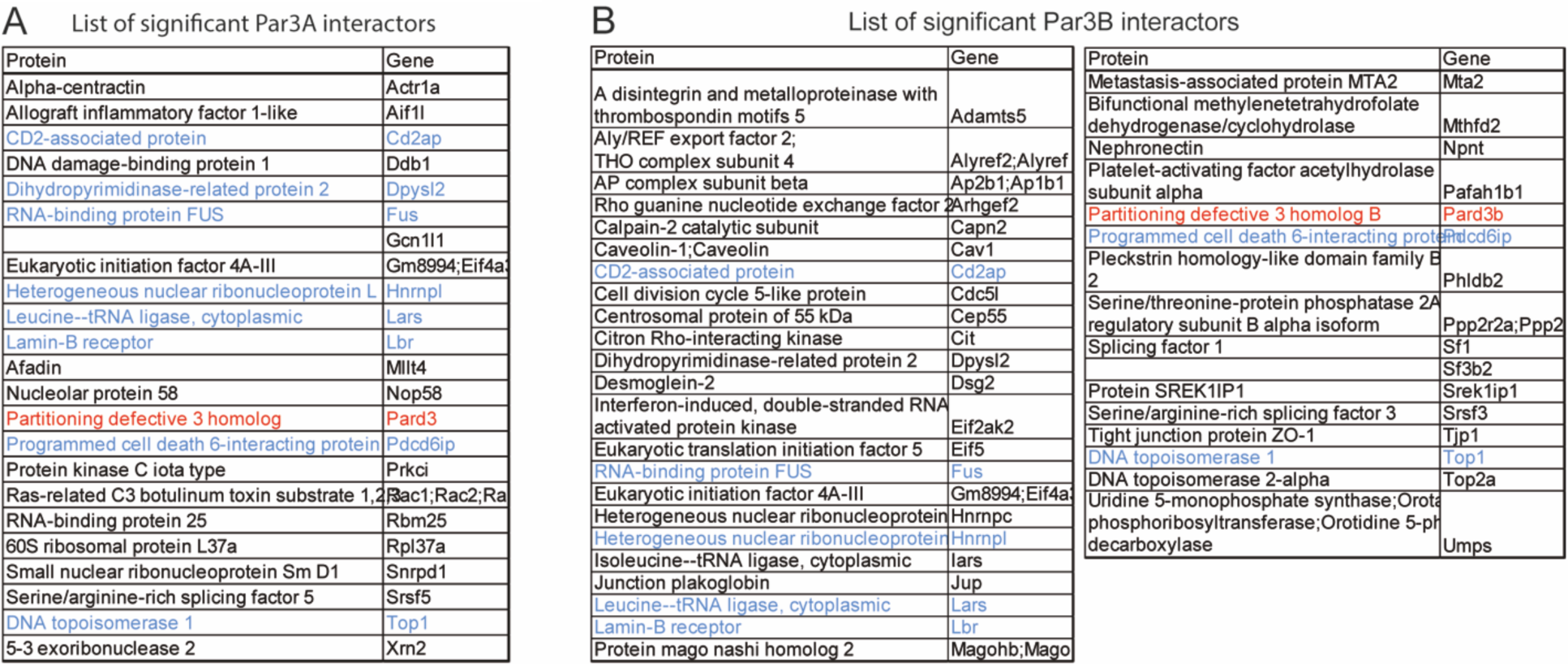
Interactome analysis of Par3A and Par3B in IMCDs. **A** List of significant Par3A interactors. Blue: shared interactors with Par3B, Red: Par3A. **B** List of significant Par3B interactors. Blue: shared interactors with Par3A, Red: Par3B.

**Supplementary Figure 8:**
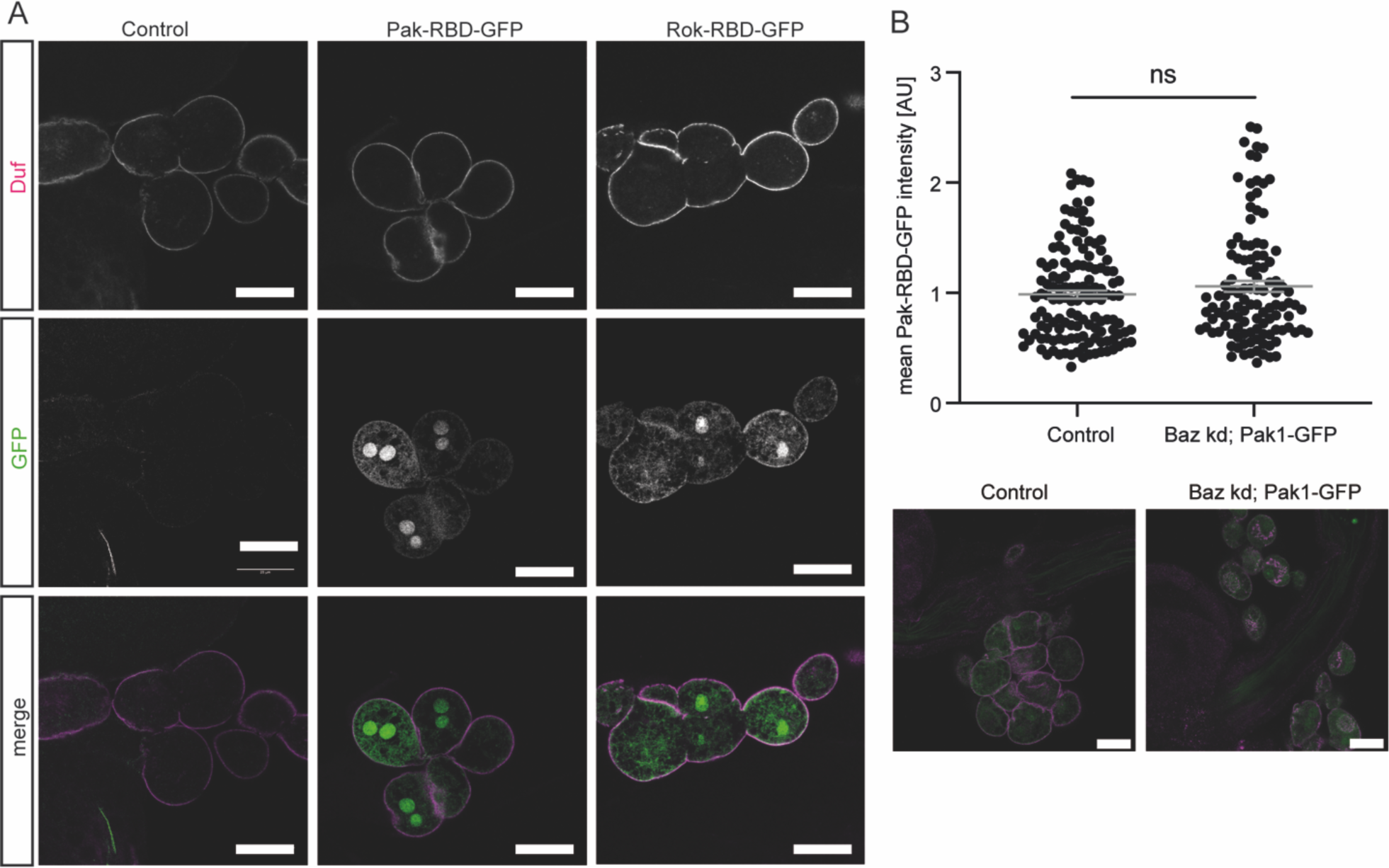
Validation of GTPase sensor flies in nephrocytes. **A** Expression of Rok-RBD-GFP and Pak-RBD-GFP was verified by confocal microscopy. Garland nephrocytes of L3 larvae were visualized using an Duf antibody and showed a clear GFP signal in both sensor fly strains. Scale bar = 25μm. Control (w;Sns-GAL4/+:UAS-Dicer/+), Pak-RBD-GFP (w; Sns-GAL4/+;UAS-Pak-RBD-GFP/UAS-Dicer), Rok-RBD-GFP (w; Sns-GAL4/+;UAS-Rok-RBD-GFP/UAS-Dicer). **B** Utilizing a Rac sensor fly strain (Pak-RBD-GFP) revealed no differences in active Rac/Cdc42 levels upon loss of Baz, when compared to control cells. (Control: w; Sns-GAL4/+;UAS-Pak-RBD-GFP/UAS-Dicer, Baz kd; Pak1-GFP: w; Sns-GAL4/UAS-Baz-RNAi;UAS-Pak-RBD-GFP/UAS-Dicer

